# Evaluating the transcriptional fidelity of cancer models

**DOI:** 10.1101/2020.03.27.012757

**Authors:** Da Peng, Rachel Gleyzer, Wen-Hsin Tai, Pavithra Kumar, Qin Bian, Bradley Issacs, Edroaldo Lummertz da Rocha, Stephanie Cai, Kathleen DiNapoli, Franklin W Huang, Patrick Cahan

## Abstract

**Background:** Cancer researchers use cell lines, patient derived xenografts, engineered mice, and tumoroids as models to investigate tumor biology and to identify therapies. The generalizability and power of a model derives from the fidelity with which it represents the tumor type under investigation, however, the extent to which this is true is often unclear. The preponderance of models and the ability to readily generate new ones has created a demand for tools that can measure the extent and ways in which cancer models resemble or diverge from native tumors.

**Methods:** We developed a machine learning based computational tool, CancerCellNet, that measures the similarity of cancer models to 22 naturally occurring tumor types and 36 subtypes, in a platform and species agnostic manner. We applied this tool to 657 cancer cell lines, 415 patient derived xenografts, 26 distinct genetically engineered mouse models, and 131 tumoroids. We validated CancerCellNet by application to independent data, and we tested several predictions with immunofluorescence.

**Results:** We have documented the cancer models with the greatest transcriptional fidelity to natural tumors, we have identified cancers underserved by adequate models, and we have found models with annotations that do not match their classification. By comparing models across modalities, we report that, on average, genetically engineered mice and tumoroids have higher transcriptional fidelity than patient derived xenografts and cell lines in four out of five tumor types. However, several patient derived xenografts and tumoroids have classification scores that are on par with native tumors, highlighting both their potential as faithful model classes and their heterogeneity.

**Conclusions:** CancerCellNet enables the rapid assessment of transcriptional fidelity of tumor models. We have made CancerCellNet available as freely downloadable software and as a web application that can be applied to new cancer models that allows for direct comparison to the cancer models evaluated here.

## INTRODUCTION

Models are widely used to investigate cancer biology and to identify potential therapeutics. Popular modeling modalities are cancer cell lines (CCLs)^1^, genetically engineered mouse models (GEMMs)^2^, patient derived xenografts (PDXs)^3^, and tumoroids^4^. These classes of models differ in the types of questions that they are designed to address. CCLs are often used to address cell intrinsic mechanistic questions^5^, GEMMs to chart progression of molecularly defined-disease^6^, and PDXs to explore patient-specific response to therapy in a physiologically relevant context^7^. More recently, tumoroids have emerged as relatively inexpensive, physiological, in vitro 3D models of tumor epithelium with applications ranging from measuring drug responsiveness to exploring tumor dependence on cancer stem cells. Models also differ in the extent to which the they represent specific aspects of a cancer type^8^. Even with this intra- and inter-class model variation, all models should represent the tumor type or subtype under investigation, and not another type of tumor, and not a non-cancerous tissue. Therefore, cancer-models should be selected not only based on the specific biological question but also based on the similarity of the model to the cancer type under investigation^9,10^.

Various methods have been proposed to determine the similarity of cancer models to their intended subjects. Domcke et al devised a ‘suitability score’ as a metric of the molecular similarity of CCLs to high grade serous ovarian carcinoma based on a heuristic weighting of copy number alterations, mutation status of several genes that distinguish ovarian cancer subtypes, and hypermutation status^11^. Other studies have taken analogous approaches by either focusing on transcriptomic or ensemble molecular profiles (e.g. transcriptomic and copy number alterations) to quantify the similarity of cell lines to tumors^12–14^. These studies were tumor-type specific, focusing on CCLs that model, for example, hepatocellular carcinoma or breast cancer. Notably, Yu et al compared the transcriptomes of CCLs to The Cancer Genome Atlas (TCGA) by correlation analysis, resulting in a panel of CCLs recommended as most representative of 22 tumor types^15^. Most recently, Najgebauer et al^16^ and Salvadores et al^17^ have developed methods to assess CCLs using molecular traits such as copy number alterations (CNA), somatic mutations, DNA methylation and transcriptomics. While all of these studies have provided valuable information, they leave two major challenges unmet. The first challenge is to determine the fidelity of GEMMs, PDXs, and tumoroids, and whether there are stark differences between these classes of models and CCLs. The other major unmet challenge is to enable the rapid assessment of new, emerging cancer models. This challenge is especially relevant now as technical barriers to generating models have been substantially lowered^18,19^, and because new models such as PDXs and tumoroids can be derived on patient-specific basis therefore should be considered a distinct entity requiring individual validation^4,20^.

To address these challenges, we developed CancerCellNet (CCN), a computational tool that uses transcriptomic data to quantitatively assess the similarity between cancer models and 22 naturally occurring tumor types and 36 subtypes in a platform- and species-agnostic manner. Here, we describe CCN’s performance, and the results of applying it to assess 657 CCLs, 415 PDXs, 26 GEMMs, and 131 tumoroids. This has allowed us to identify the most faithful models currently available, to document cancers underserved by adequate models, and to find models with inaccurate tumor type annotation. Moreover, because CCN is open-source and easy to use, it can be readily applied to newly generated cancer models as a means to assess their fidelity.

## RESULTS

### CancerCellNet classifies samples accurately across species and technologies

Previously, we had developed a computational tool using the Random Forest classification method to measure the similarity of engineered cell populations to their *in vivo* counterparts based on transcriptional profiles^21,22^. More recently, we elaborated on this approach to allow for classification of single cell RNA-seq data in a manner that allows for cross-platform and cross-species analysis^23^. Here, we used an analogous approach to build a platform that would allow us to quantitatively compare cancer models to naturally occurring patient tumors (**Fig 1A**). In brief, we used TCGA RNA-seq expression data from 22 solid tumor types to train a top-pair multi-class Random forest classifier (**Fig 1B**). We combined training data from Rectal Adenocarcinoma (READ) and Colon Adenocarcinoma (COAD) into one COAD_READ category because READ and COAD are considered to be virtually indistinguishable at a molecular level^24^. We included an ‘Unknown’ category trained using randomly shuffled gene-pair profiles generated from the training data of 22 tumor types to identify query samples that are not reflective of any of the training data. To estimate the performance of CCN and how it is impacted by parameter variation, we performed a parameter sweep with a 5-fold 2/3 cross-validation strategy (i.e. 2/3 of the data sampled across each cancer type was used to train, 1/3 was used to validate) (**Fig 1C**). The performance of CCN, as measured by the mean area under the precision recall curve (AUPRC), did not fall below 0.945 and remained relatively stable across parameter sets (**Supp Fig 1A**). The optimal parameters resulted in 1,979 features. The mean AUPRCs exceeded 0.95 in most tumor types with this optimal parameter set (**Fig 1D, Supp Fig 1B**). The AUPRCs of CCN applied to independent data RNA-Seq data from 725 tumors across five tumor types from the International Cancer Genome Consortium (ICGC)^25^ ranged from 0.93 to 0.99, supporting the notion that the platform is able to accurately classify tumor samples from diverse sources (**Fig 1E**).

**Fig. 1.**
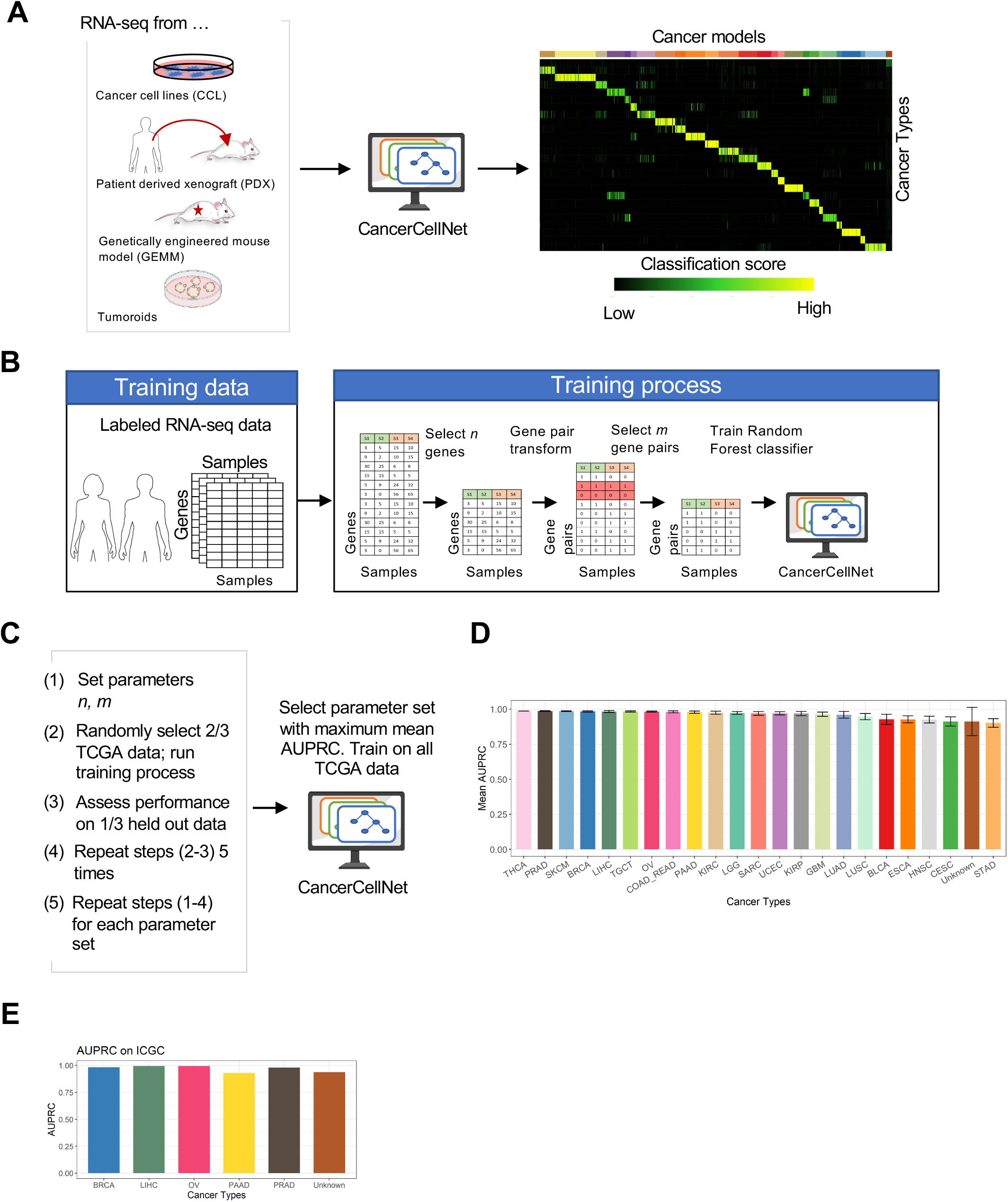
CancerCellNet (CCN) workflow, training, and performance. **(A)** Schematic of CCN usage. CCN was designed to assess and compare the expression profiles of cancer models such as CCLs, PDXs, GEMMs, and tumoroids with native patient tumors. To use trained classifier, CCN inputs the query samples (e.g. expression profiles from CCLs, PDXs, GEMMs, tumoroids) and generates a classification profile for the query samples. The column names of the classification heatmap represent sample annotation and the row names of the classification heatmap represent different cancer types. Each grid is colored from black to yellow representing the lowest classification score (e.g. 0) to highest classification score (e.g. 1). **(B)** Schematic of CCN training process. CCN uses patient tumor expression profiles of 22 different cancer types from TCGA as training data. First, CCN identifies *n* genes that are upregulated, *n* that are downregulated, and *n* that are relatively invariant in each tumor type versus all of the others. Then, CCN performs a pair transform on these genes and subsequently selects the most discriminative set of *m* gene pairs for each cancer type as features (or predictors) for the Random forest classifier. Lastly, CCN trains a multi-class Random Forest classifier using gene-pair transformed training data. **(C)** Parameter optimization strategy. 5 cross-validations of each parameter set in which 2/3 of TCGA data was used to train and 1/3 to validate was used search for the values of *n* and *m* that maximized performance of the classifier as measured by area under the precision recall curve (AUPRC). **(D)** Mean and standard deviation of classifiers based on 50 cross-validations with the optimal parameter set. **(E)** AUPRC of the final CCN classifier when applied to independent patient tumor data from ICGC.

As one of the central aims of our study is to compare distinct cancer models, including GEMMs, our method needed to be able to classify samples from mouse and human samples equivalently. We used the Top-Pair transform^23^ to achieve this and we tested the feasibility of this approach by assessing the performance of a normal (i.e. non-tumor) cell and tissue classifier trained on human data as applied to mouse samples. Consistent with prior applications^23^, we found that the cross-species classifier performed well, achieving mean AUPRC of 0.97 when applied to mouse data (**Supp Fig 1C**).

To evaluate cancer models at a finer resolution, we also developed an approach to perform tumor subtype classifications (**Supp Fig 1D**). We constructed 11 different cancer subtype classifiers based on the availability of expression or histological subtype information^24,26–36^. We also included non-cancerous, normal tissues as categories for several subtype classifiers when sufficient data was available: breast invasive carcinoma (BRCA), COAD_READ, head and neck squamous cell carcinoma (HNSC), kidney renal clear cell carcinoma (KIRC) and uterine corpus endometrial carcinoma (UCEC). The 11 subtype classifiers all achieved high overall average AUPRs ranging from 0.80 to 0.99 (**Supp Fig 1E**).

### Fidelity of cancer cell lines

Having validated the performance of CCN, we then used it to determine the fidelity of CCLs. We mined RNA-seq expression data of 657 different cell lines across 20 cancer types from the Cancer Cell Line Encyclopedia (CCLE) and applied CCN to them, finding a wide classification range for cell lines of each tumor type (**Fig 2A, Supp Tab 1**). To verify the classification results, we applied CCN to expression profiles from CCLE generated through microarray expression profiling^37^. To ensure that CCN would function on microarray data, we first tested it by applying a CCN classifier created to test microarray data to 720 expression profiles of 12 tumor types. The cross-platform CCN classifier performed well, based on the comparison to study-provided annotation, achieving a mean AUPRC of 0.91 (**Supp Fig 2A**). Next, we applied this cross-platform classifier to microarray expression profiles from CCLE (**Supp Fig 2B**). From the classification results of 571 cell lines that have both RNA-seq and microarray expression profiles, we found a strong overall positive association between the classification scores from RNA-seq and those from microarray (**Supp Fig 2C**). This comparison supports the notion that the classification scores for each cell line are not artifacts of profiling methodology. Moreover, this comparison shows that the scores are consistent between the times that the cell lines were first assayed by microarray expression profiling in 2012 and by RNA-Seq in 2019. We also observed high level of correlation between our analysis and the analysis done by Yu et al^15^(**Supp Fig 2D**), further validating the robustness of the CCN results.

**Fig. 2.**
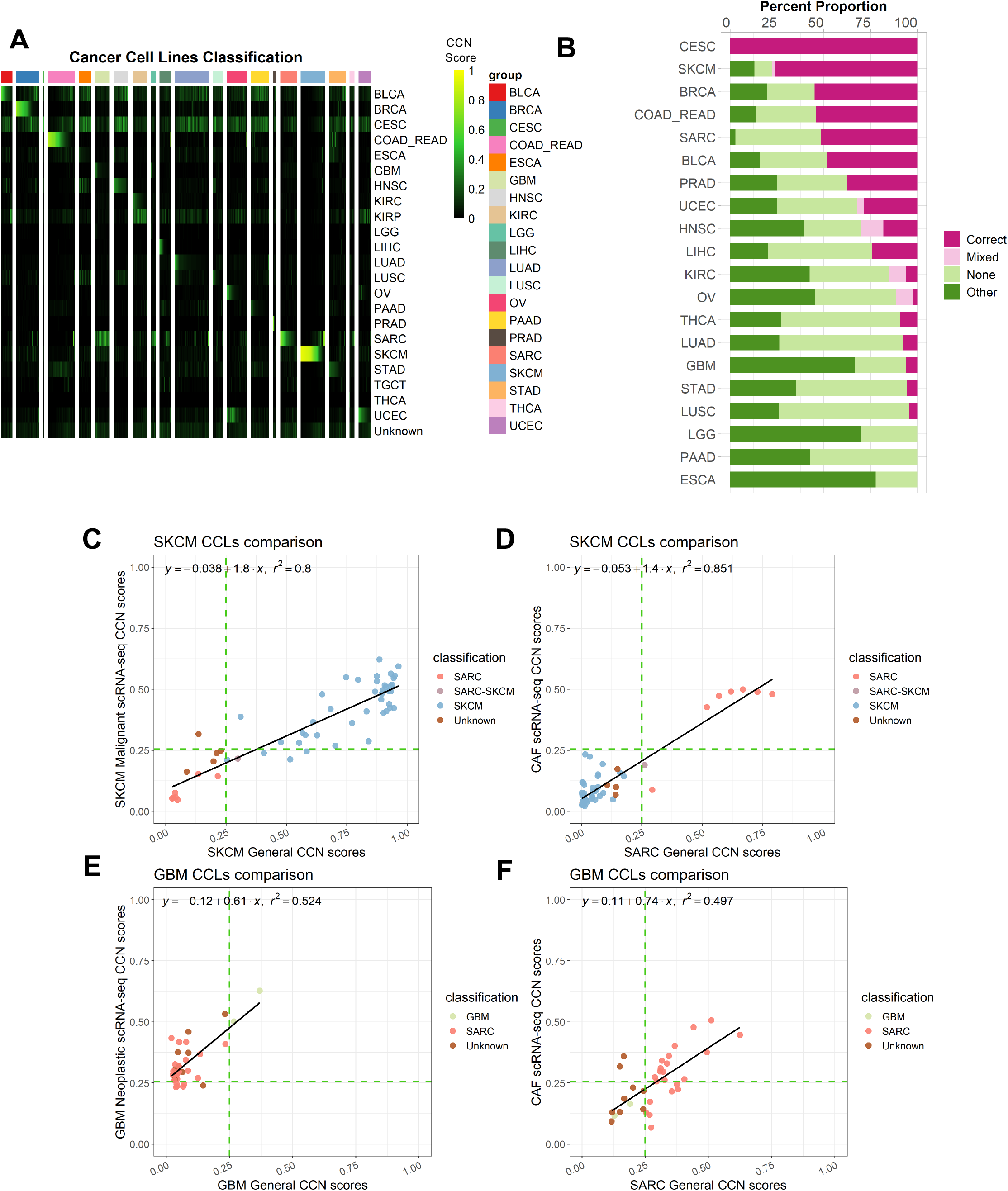
Evaluation of cancer cell lines. **(A)** General classification heatmap of CCLs extracted from CCLE. Column annotations of the heatmap represent the labelled cancer category of the CCLs given by CCLE and the row names of the heatmap represent different cancer categories. CCLs’ general classification profiles are categorized into 4 categories: correct (red), correct mixed (pink), no classification (light green) and other classification (dark green) based on the decision threshold of 0.25. **(B)** Bar plot represents the proportion of each classification category in CCLs across cancer types ordered from the cancer types with the highest proportion of correct and correct mixed CCLs to lowest proportion. **(C)** Comparison between SKCM general CCN scores from bulk RNA-seq classifier and SKCM malignant CCN scores from scRNA-seq classifier for SKCM CCLs. **(D)** Comparison between SARC general CCN scores from bulk RNA-seq classifier and CAF CCN scores from scRNA-seq classifier for SKCM CCLs. **(E)** Comparison between GBM general CCN scores from bulk RNA-seq classifier and GBM neoplastic CCN scores from scRNA-seq classifier for GBM CCLs. **(F)** Comparison between SARC general CCN scores and CAF CCN scores from scRNA-seq classifier for GBM CCLs. The green lines indicate the decision threshold for scRNA-seq classifier and general classifier.

Next, we assessed the extent to which CCN classifications agreed with their nominal tumor type of origin, which entailed translating quantitative CCN scores to classification labels. To achieve this, we selected a decision threshold that maximized the Macro F1 measure, harmonic mean of precision and recall, across 50 cross validations. Then, we annotated cell lines based their CCN score profile as follows. Cell lines with CCN scores > threshold for the tumor type of origin were annotated as ‘correct’. Cell lines with CCN scores > threshold in the tumor type of origin and at least one other tumor type were annotated as ‘mixed’. Cell lines with CCN scores > threshold for tumor types other than that of the cell line’s origin were annotated as ‘other’. Cell lines that did not receive a CCN score > threshold for any tumor type were annotated as ‘none’ (**Fig 2B**). We found that majority of cell lines originally annotated as Breast invasive carcinoma (BRCA), Cervical squamous cell carcinoma and endocervical adenocarcinoma (CESC), Skin Cutaneous Melanoma (SKCM), Colorectal Cancer (COAD_READ) and Sarcoma (SARC) fell into the ‘correct’ category (**Fig 2B**). On the other hand, no Esophageal carcinoma (ESCA), Pancreatic adenocarcinoma (PAAD) or Brain Lower Grade Glioma (LGG) were classified as ‘correct’, demonstrating the need for more transcriptionally faithful cell lines that model those general cancer types.

There are several possible explanations for cell lines not receiving a ‘correct’ classification. One possibility is that the sample was incorrectly labeled in the study from which we harvested the expression data. Consistent with this explanation, we found that colorectal cancer line NCI-H684^38,39^, a cell line labelled as liver hepatocellular carcinoma (LIHC) by CCLE, was classified strongly as COAD_READ (**Supp Tab 1**). Another possibility to explain low CCN score is that cell lines were derived from subtypes of tumors that are not well-represented in TCGA. To explore this hypothesis, we first performed tumor subtype classification on CCLs from 11 tumor types for which we had trained subtype classifiers (**Supp Tab 2**). We reasoned that if a cell was a good model for a rarer subtype, then it would receive a poor general classification but a high classification for the subtype that it models well. Therefore, we counted the number of lines that fit this pattern. We found that of the 188 lines with no general classification, 25 (13%) were classified as a specific subtype, suggesting that derivation from rare subtypes is not the major contributor to the poor overall fidelity of CCLs.

Another potential contributor to low scoring cell lines is intra-tumor stromal and immune cell impurity in the training data. If impurity were a confounder of CCN scoring, then we would expect a strong positive correlation between mean purity and mean CCN classification scores of CCLs per general tumor type. However, the Pearson correlation coefficient between the mean purity of general tumor type and mean CCN classification scores of CCLs in the corresponding general tumor type was low (0.14), suggesting that tumor purity is not a major contributor to the low CCN scores across CCLs (**Supp Fig 2E**).

### Comparison of SKCM and GBM CCLs to scRNA-seq

To more directly assess the impact of intra-tumor heterogeneity in the training data on evaluating cell lines, we constructed a classifier using cell types found in human melanoma and glioblastoma scRNA-seq data^40,41^. Previously, we have demonstrated the feasibility of using our classification approach on scRNA-seq data^23^. Our scRNA-seq classifier achieved a high average AUPRC (0.95) when applied to held-out data and high mean AUPRC (0.99) when applied to few purified bulk testing samples (**Supp Fig 3A-B**). Comparing the CCN score from bulk RNA-seq general classifier and scRNA-seq classifier, we observed a high level of correlation (Pearson correlation of 0.89) between the SKCM CCN classification scores and scRNA-seq SKCM malignant CCN classification scores for SKCM cell lines (**Fig 2C, Supp Fig 3C**). Of the 41 SKCM cell lines that were classified as SKCM by the bulk classifier, 37 were also classified as SKCM malignant cells by the scRNA-seq classifier. Interestingly, we also observed a high correlation between the SARC CCN classification score and scRNA-seq cancer associated fibroblast (CAF) CCN classification scores (Pearson correlation of 0.92). Six of the seven SKCM cell lines that had been classified as exclusively SARC by CCN were classified as CAF by the scRNA-seq classifier (**Fig 2D, Supp Fig 3C**), which suggests the possibility that these cell lines were derived from CAF or other mesenchymal populations, or that they have acquired a mesenchymal character through their derivation. The high level of agreement between scRNA-seq and bulk RNA-seq classification results shows that heterogeneity in the training data of general CCN classifier has little impact in the classification of SKCM cell lines.

In contrast, we observed a weaker correlation between GBM CCN classification scores and scRNA-seq GBM neoplastic CCN classification scores (Pearson correlation of 0.72) for GBM cell lines (**Fig 2E, Supp Fig 3D**). Of the 31 GBM lines that were not classified as GBM with CCN, 25 were classified as GBM neoplastic cells with the scRNA-seq classifier. Among the 22 GBM lines that were classified as SARC with CCN, 15 cell lines were classified as CAF (**Fig 2F**), 10 which were classified as both GBM neoplastic and CAF in the scRNA-seq classifier. Similar to the situation with SKCM lines that classify as CAF, this result is consistent with the possibility that some GBM lines classified as SARC by CCN could be derived from mesenchymal subtypes exhibiting both strong mesenchymal signatures and glioblastoma signatures or that they have acquired a mesenchymal character through their derivation. The lower level of agreement between scRNA-seq and bulk RNA-seq classification results for GBM models suggests that the heterogeneity of glioblastomas^42^ can impact the classification of GBM cell lines, and that the use of scRNA-seq classifier can resolve this deficiency.

### Immunofluorescence confirmation of CCN predictions

To experimentally explore some of our computational analyses, we performed immunofluorescence on three cell lines that were not classified as their labelled categories: the ovarian cancer line SK-OV-3 had a high UCEC CCN score (0.246), the ovarian cancer line A2780 had a high Testicular Germ Cell Tumors (TGCT) CCN score (0.327), and the prostate cancer line PC-3 had a high bladder cancer (BLCA) score (0.307) (**Supp Tab 1**). We reasoned that if SK-OV-3, A2780 and PC-3 were classified most strongly as UCEC, TGCT and BLCA, respectively, then they would express proteins that are indicative of these cancer types.

First, we measured the expression of the uterine-associated transcription factor HOXB6^43,44^, and the UCEC serous ovarian tumor biomarker WT1^45^ in SK-OV-3, in the OV cell line Caov-4, and in the UCEC cell line HEC-59. We chose Caov-4 as our positive control for OV biomarker expression because it was determined by our analysis and others^11,15^ to be a good model of OV. Likewise, we chose HEC-59 to be a positive control for UCEC. We found that SK-OV-3 has a small percentage (5%) of cells that expressed the uterine marker HOXB6 and a large proportion (73%) of cells that expressed WT1 (**Fig 3A**). In contrast, no Caov-4 cells expressed HOXB6, whereas 85% of cells expressed WT1. This suggests that SK-OV-3 exhibits both biomarkers of ovarian tumor and uterine tissue. From our computational analysis and experimental validation, SK-OV-3 is most likely an endometrioid subtype of ovarian cancer. This result is also consistent with prior classification of SK-OV-3^46^, and the fact that SK-OV-3 lacks p53 mutations, which is prevalent in high-grade serous ovarian cancer^47^, and it harbors an endometrioid-associated mutation in ARID1A^11,46,48^. Next, we measured the expression of markers of OV and germ cell cancers (LIN28A^49^) in the OV-annotated cell line A2780, which received a high TCGT CCN score. We found that 54% of A2780 cells expressed LIN28A whereas it was not detected in Caov-4 (**Fig 3B**). The OV marker WT1 was also expressed in fewer A2780 cells as compared to Caov-4 (48% vs 85%), which suggests that A2780 could be a germ cell derived ovarian tumor. Taken together, our results suggest that SK-OV-3 and A2780 could represent OV subtypes of that are not well represented in TCGA training data, which resulted in a low OV score and higher CCN score in other categories.

**Fig. 3.**
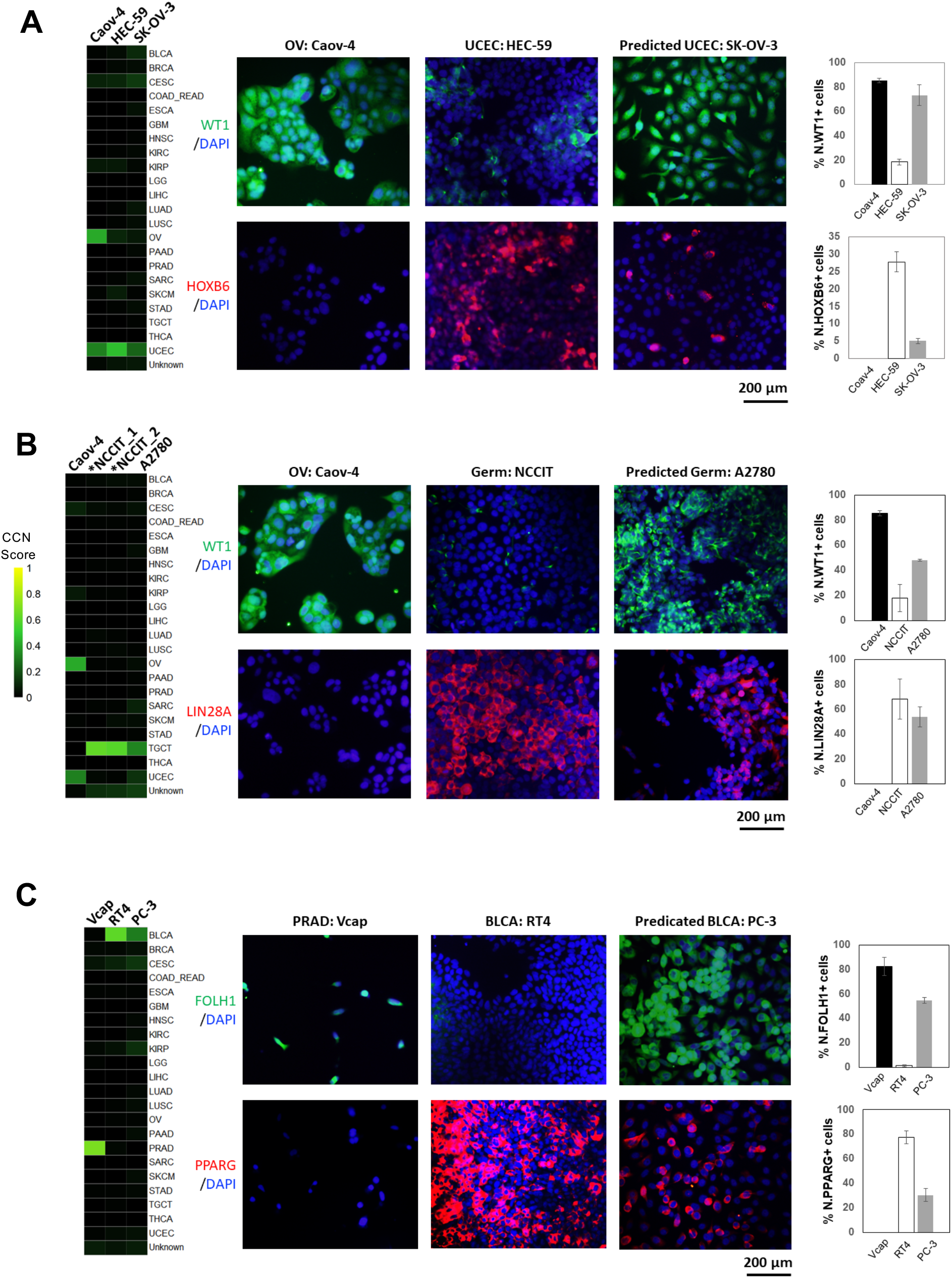
Immunofluorescence of selected cell lines. **(A)** Classification profiles (left) and IF expression (middle) of Caov-4 (OV positive control), HEC-59 (UCEC positive control) and SK-OV-3 for WT1 (OV biomarker) and HOXB6 (uterine biomarker). The bar plots quantify the average percentage of positive cells for WT1 (top-right) and HOXB6 (bottom-right). **(B)** Classification profiles (left) and IF expression (middle) of Caov-4, NCCIT (germ cell tumor positive control) and A2780 for WT1 and LIN28A (germ cell tumor biomarker). Classification of NCCIT were performed using RNA-seq profiles of WT control NCCIT duplicate from Grow et al^91^. The bar plots quantify the average percentage of positive cells for WT1 (top-right) and LIN28A (bottom-right). **(C)** Classification profiles (left) and IF expression (middle) of Vcap (PRAD positive control), RT4 (BLCA positive control) and PC-3 for FOLH1 (prostate biomarker) and PPARG (urothelial biomarker). The bar plots quantify the average percentage of positive cells for FOLH1 (top-right) and PPARG (bottom-right).

Lastly, we examined PC-3, annotated as a PRAD cell line but classified to be most similar to BLCA. We found that 30% of the PC-3 cells expressed PPARG, a contributor to urothelial differentiation^50^ that is not detected in the PRAD Vcap cell line but is highly expressed in the BLCA RT4 cell line (**Fig 3C**). PC-3 cells also expressed the PRAD biomarker FOLH1^51^ suggesting that PC-3 has an PRAD origin and gained urothelial or luminal characteristics through the derivation process. In short, our limited experimental data support the CCN classification results.

### Subtype classification of cancer cell lines

Next, we explored the subtype classification of CCLs from three general tumor types in more depth. We focused our subtype visualization (**Fig 4A-C**) on CCL models with general CCN score above 0.1 in their nominal cancer type as this allowed us to analyze those models that fell below the general threshold but were classified as a specific sub-type (**Supp Tab 1-2**).

**Fig. 4.**
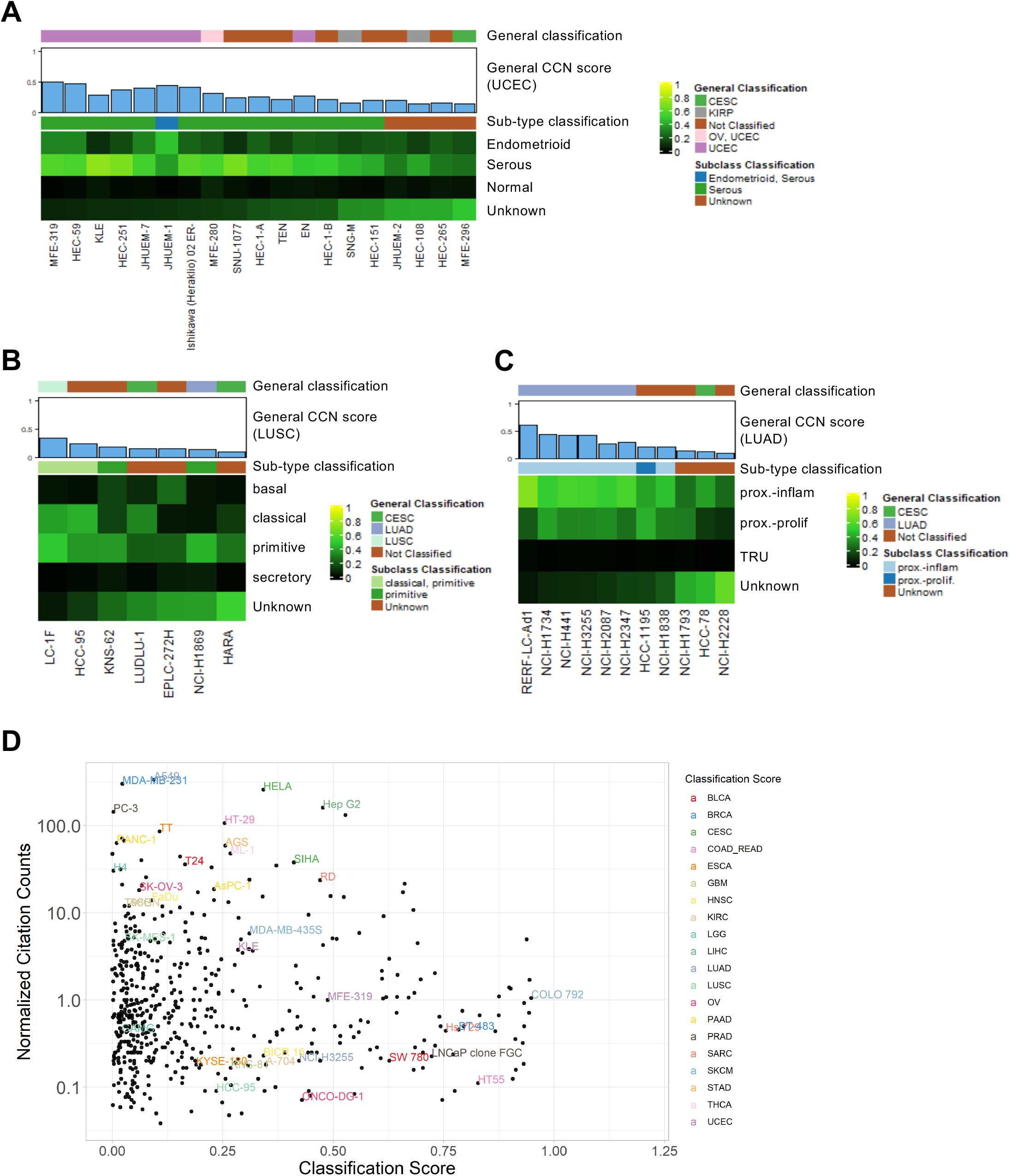
Subtype classification of CCLs and CCL prevalence. The heatmap visualizations represent subtype classification of **(A)** UCEC CCLs**, (B)** LUSC CCLs and **(C)** LUAD CCLs. Only samples with CCN scores > 0.1 in their nominal tumor type are displayed. **(D)** Comparison of normalized citation counts and general CCN classification scores of CCLs. Labelled cell lines either have the highest CCN classification score in their labelled cancer category or highest normalized citation count. Each citation count was normalized by number of years since first documented on PubMed.

Focusing first on UCEC, the histologically defined subtypes of UCEC, endometrioid and serous, differ in prevalence, molecular properties, prognosis, and treatment. For instance, the endometrioid subtype, which accounts for approximately 80% of uterine cancers, retains estrogen receptor and progesterone receptor status and is responsive towards progestin therapy^52,53^. Serous, a more aggressive subtype, is characterized by the loss of estrogen and progesterone receptor and is not responsive to progestin therapy^52,53^. CCN classified the majority of the UCEC cell lines as serous except for JHUEM-1 which is classified as mixed, with similarities to both endometrioid and serous (**Fig 4A**). The preponderance CCLE lines of serous versus endometroid character may be due to properties of serous cancer cells that promote their *in vitro* propagation, such as upregulation of cell adhesion transcriptional programs^54^. Some of our subtype classification results are consistent with prior observations. For example, HEC-1A, HEC-1B, and KLE were previously characterized as type II endometrial cancer, which includes a serous histological subtype^55^. On the other hand, our subtype classification results contradict prior observations in at least one case. For instance, the Ishikawa cell line was derived from type I endometrial cancer (endometrioid histological subtype)^55,56^, however CCN classified a derivative of this line, Ishikawa 02 ER-, as serous. The high serous CCN score could result from a shift in phenotype of the line concomitant with its loss of estrogen receptor (ER) as this is a distinguishing feature of type II endometrial cancer (serous histological subtype)^52^. Taken together, these results indicate a need for more endometroid-like CCLs.

Next, we examined the subtype classification of Lung Squamous Cell Carcinoma (LUSC) and Lung adenocarcinoma (LUAD) cell lines (Fig 4B-C). All the LUSC lines with at least one subtype classification had an underlying primitive subtype classification. This is consistent either with the ease of deriving lines from tumors with a primitive character, or with a process by which cell line derivation promotes similarity to more primitive subtype, which is marked by increased cellular proliferation^28^. Some of our results are consistent with prior reports that have investigated the resemblance of some lines to LUSC subtypes. For example, HCC-95, previously been characterized as classical^28,57^, had a maximum CCN score in the classical subtype (0.429). Similarly, LUDLU-1 and EPLC-272H, previously reported as classical^57^ and basal^57^ respectively, had maximal tumor subtype CCN scores for these sub-types (0.323 and 0.256) (**Fig 4B, Supp Tab 2**) despite classified as Unknown. Lastly, the LUAD cell lines that were classified as a subtype were either classified as proximal inflammation or proximal proliferation (**Fig 4C**). RERF-LC-Ad1 had the highest general classification score and the highest proximal inflammation subtype classification score. Taken together, these subtype classification results have revealed an absence of cell lines models for basal and secretory LUSC, and for the Terminal respiratory unit (TRU) LUAD subtype.

### Cancer cell lines’ popularity and transcriptional fidelity

Finally, we sought to measure the extent to which cell line transcriptional fidelity related to model prevalence. We used the number of papers in which a model was mentioned, normalized by the number of years since the cell line was documented, as a rough approximation of model prevalence. To explore this relationship, we plotted the normalized citation count versus general classification score, labeling the highest cited and highest classified cell lines from each general tumor type (**Fig 4D**). For most of the general tumor types, the highest cited cell line is not the highest classified cell line except for Hep G2, AGS and ML-1, representing liver hepatocellular carcinoma (LIHC), stomach adenocarcinoma (STAD), and thyroid carcinoma (THCA), respectively. On the other hand, the general scores of the highest cited cell lines representing BLCA (T24), BRCA (MDA-MB-231), and PRAD (PC-3) fall below the classification threshold of 0.25. Notably, each of these tumor types have other lines with scores exceeding 0.5, which should be considered as more faithful transcriptional models when selecting lines for a study (**Supp Tab 1 and http://www.cahanlab.org/resources/cancerCellNet_results/**).

### Evaluation of patient derived xenografts

Next, we sought to evaluate a more recent class of cancer models: PDX. To do so, we subjected the RNA-seq expression profiles of 415 PDX models from 13 different types of cancer types generated previously^20^ to CCN. Similar to the results of CCLs, the PDXs exhibited a wide range of classification scores (**Fig 5A, Supp Tab 3**). By categorizing the CCN scores of PDX based on the proportion of samples associated with each tumor type that were correctly classified, we found that SARC, SKCM, COAD_READ and BRCA have higher proportion of correctly classified PDX than those of other cancer categories (**Fig 5B**). In contrast to CCLs, we found a higher proportion of correctly classified PDX in STAD, PAAD and KIRC (**Fig 5B**). However, similar to CCLs, no ESCA PDXs were classified as such. This held true when we performed subtype classification on PDX samples: none of the PDX in ESCA were classified as any of the ESCA subtypes (**Supp Tab 4**). UCEC PDXs had both endometrioid subtypes, serous subtypes, and mixed subtypes, which provided a broader representation than CCLs (**Fig 5C**). Several LUSC PDXs that were classified as a subtype were also classified as Head and Neck squamous cell carcinoma (HNSC) or mix HNSC and LUSC (**Fig 5D**). This could be due to the similarity in expression profiles of basal and classical subtypes of HNSC and LUSC^28,58^, which is consistent with the observation that these PDXs were also subtyped as classical. No LUSC PDXs were classified as the secretory subtype. In contrast to LUAD CCLs, four of the five LUAD PDXs with a discernible sub-type were classified as proximal inflammatory (**Fig 5E**). On the other hand, similar to the CCLs, there were no TRU subtypes in the LUAD PDX cohort. In summary, we found that while individual PDXs can reach extremely high transcriptional fidelity to both general tumor types and subtypes, many PDXs were not classified as the general tumor type from which they originated.

**Fig. 5.**
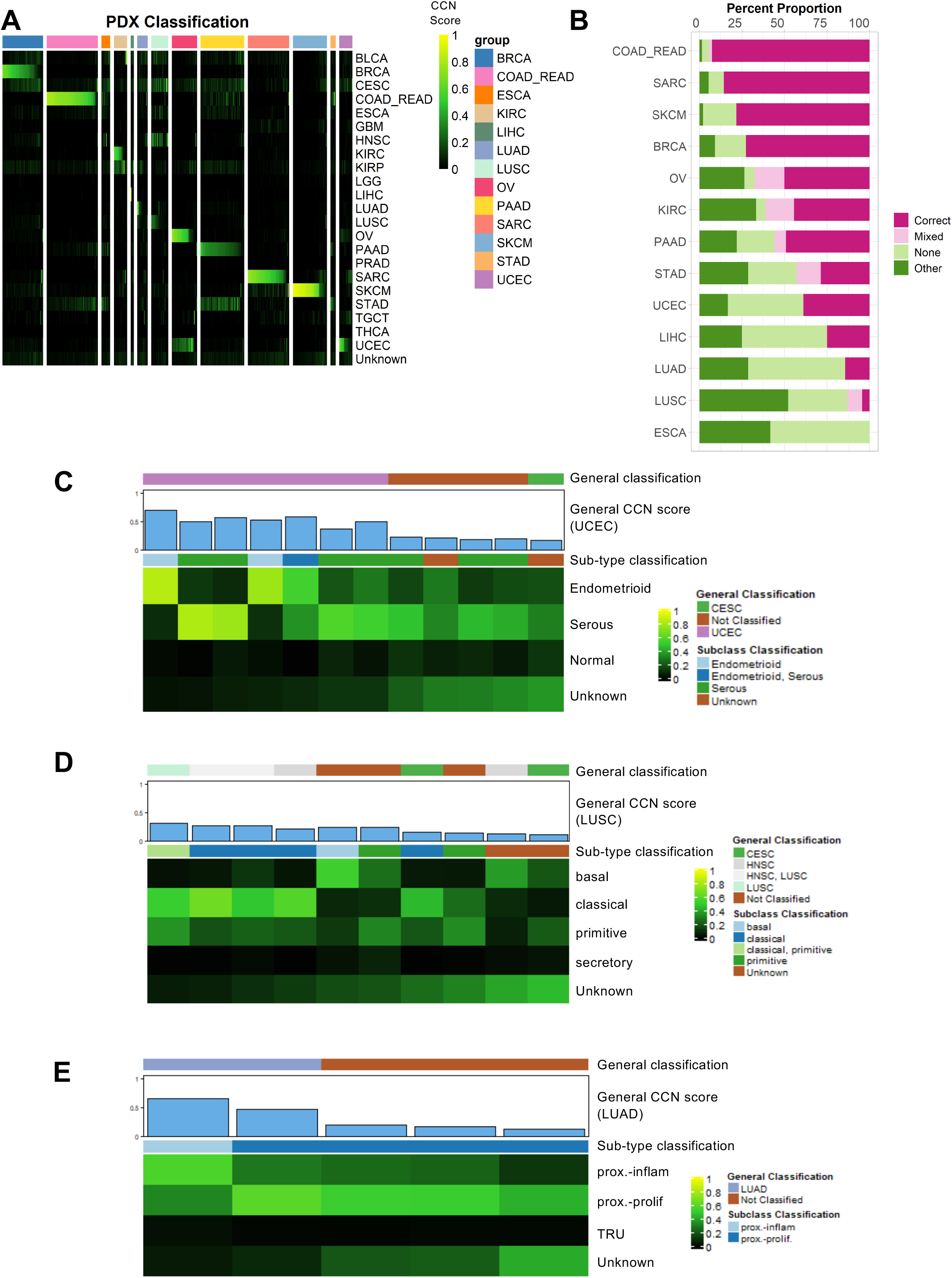
Evaluation of patient derived xenografts. **(A)** General classification heatmap of PDXs. Column annotations represent annotated cancer type of the PDXs, and row names represent cancer categories. **(B)** Proportion of classification categories in PDXs across cancer types is visualized in the bar plot and ordered from the cancer type with highest proportion of correct and mixed correct classified PDXs to the lowest. Subtype classification heatmaps of **(C)** UCEC PDXs, **(D)** LUSC PDXs and **(E)** LUAD PDXs. Only samples with CCN scores > 0.1 in their nominal tumor type are displayed.

### Evaluation of GEMMs

Next, we used CCN to evaluate GEMMs of six general tumor types from nine studies for which expression data was publicly available^59–67^. As was true for CCLs and PDXs, GEMMs also had a wide range of CCN scores (**Fig 6A, Supp Tab 5**). We next categorized the CCN scores based on the proportion of samples associated with each tumor type that were correctly classified (**Fig 6B**). In contrast to LGG CCLs, LGG GEMMs, generated by Nf1 mutations expressed in different neural progenitors in combination with Pten deletion^66^, consistently were classified as LGG (**Fig 6A-B**). The GEMM dataset included multiple replicates per model, which allowed us to examine intra-GEMM variability. Both at the level of CCN score and at the level of categorization, GEMMs were invariant. For example, replicates of UCEC GEMMs driven by Prg(cre/+)Pten(lox/lox) received almost identical general CCN scores (**Fig 6C, Supp Tab 6**). GEMMs sharing genotypes across studies, such as LUAD GEMMs driven by Kras mutation and loss of p53^59,65,67^, also received similar general and subtype classification scores (**Fig 6A,B,E**).

**Fig. 6.**
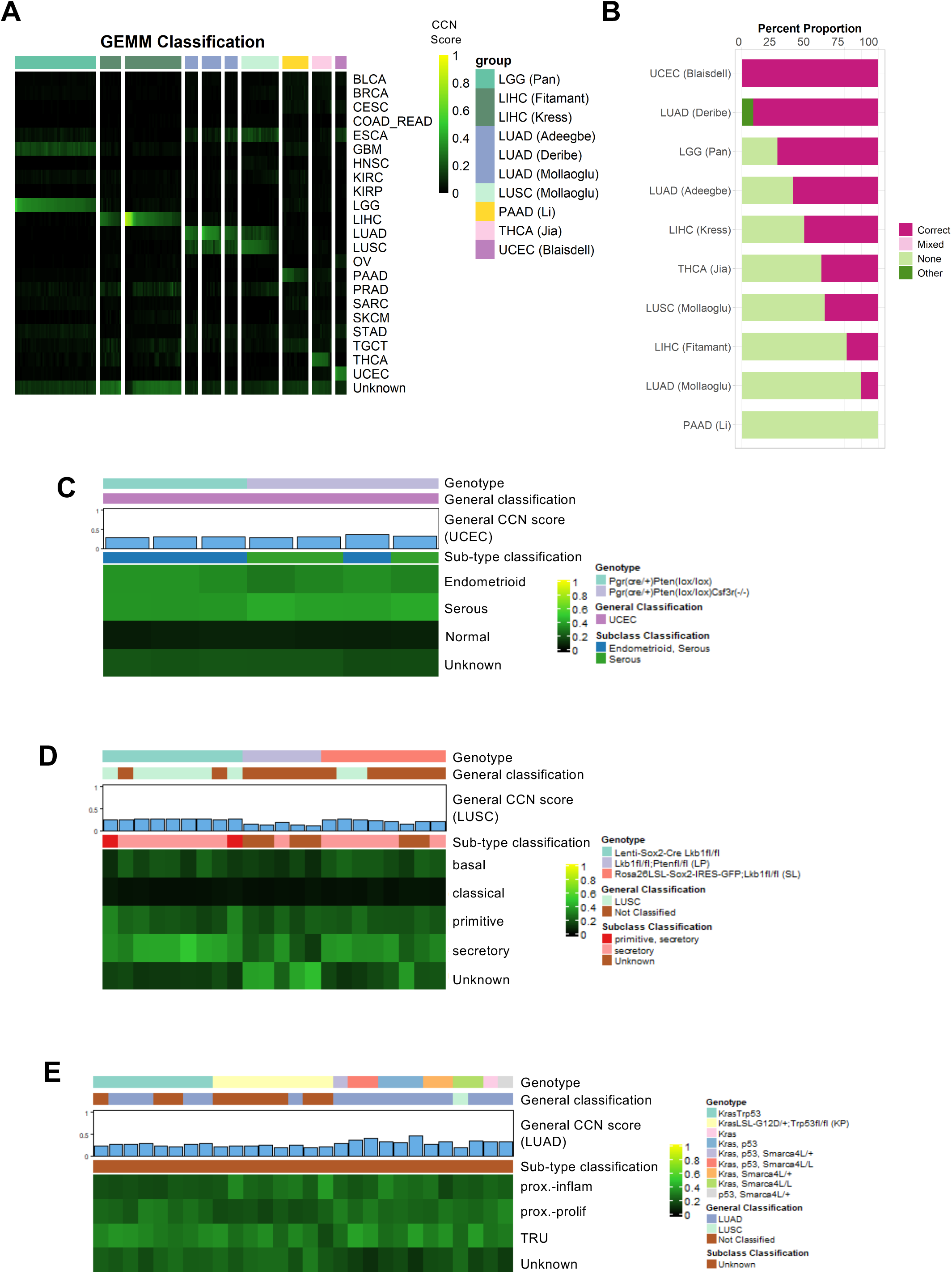
Evaluation of genetically engineered mouse models. **(A)** General classification heatmap of GEMMs. Column annotations represent annotated cancer type of the GEMMs, and row names represent cancer categories. **(B)** Proportion of classification categories in GEMMs across cancer types is visualized in the bar plot and ordered from the cancer type with highest proportion of correct and mixed correct classified GEMMs to the lowest. Subtype classification heatmap of **(C)** UCEC GEMMs, **(D)** LUSC GEMMs and **(E)** LUAD GEMMs. Only samples with CCN scores > 0.1 in their nominal tumor type are displayed.

Next, we explored the extent to which genotype impacted subtype classification in UCEC, LUSC, and LUAD. Prg(cre/+)Pten(lox/lox) GEMMs had a mixed subtype classification of both serous and endometrioid, consistent with the fact that Pten loss occurs in both subtypes (albeit more frequently in endometrioid). We also analyzed Prg(cre/+)Pten(lox/lox)Csf3r-/-GEMMs. Polymorphonuclear neutrophils (PMNs), which play anti-tumor roles in endometrioid cancer progression, are depleted in these animals. Interestingly, Prg(cre/+)Pten(lox/lox)Csf3r-/-GEMMs had a serous subtype classification, which could be explained by differences in PMN involvement in endometrioid versus serous uterine tumor development that are reflected in the respective transcriptomes of the TCGA UCEC training data. We note that the tumor cells were sorted prior to RNA-seq and thus the shift in subtype classification is not due to contamination of GEMMs with non-tumor components. In short, this analysis supports the argument that tumor-cell extrinsic factors, in this case a reduction in anti-tumor PMNs, can shift the transcriptome of a GEMM so that it more closely resembles a serous rather than endometrioid subtype.

The LUSC GEMMs that we analyzed were Lkb1^fl/fl^ and they either overexpressed of Sox2 (via two distinct mechanisms) or were also Pten^fl/fl^^65^. We note that the eight lenti-Sox2-Cre-infected;Lkb1^fl/fl^ and Rosa26LSL-Sox2-IRES-GFP;Lkb1^fl/fl^ samples that classified as ‘Unknown’ had LUSC CCN scores only modestly lower than the decision threshold (**Fig 6D**) (mean CCN score = 0.217). Thirteen out of the 17 of the Sox2 GEMMs classified as the secretory subtype of LUSC. The consistency is not surprising given both models overexpress Sox2 and lose Lkb1. On the other hand, the Lkb1^fl/fl^;Pten^fl/fl^ GEMMs had substantially lower general LUSC CCN scores and our subtype classification indicated that this GEMM was mostly classified as ‘Unknown’, in contrast to prior reports suggesting that it is most similar to a basal subtype^68^. None of the three LUSC GEMMs have strong classical CCN scores. Most of the LUAD GEMMs, which were generated using various combinations of activating Kras mutation, loss of Trp53, and loss of Smarca4L^59,65,67^, were correctly classified (**Fig 6E**). Those that were not classified have modestly lower CCN score than the decision threshold (mean CCN score = 0.214). There were no substantial differences in general or subtype classification across driver genotypes. Although the sub-type of all LUAD GEMMs was ‘Unknown’, the subtypes tended to have a mixture of high CCN proximal proliferation, proximal inflammation and TRU scores. Taken together, this analysis suggests that there is a degree of similarity, and perhaps plasticity between the primitive and secretory (but not basal or classical) subtypes of LUSC. On the other hand, while the LUAD GEMMs classify strongly as LUAD, they do not have strong particular subtype classification -- a result that does not vary by genotype.

### Evaluation of Tumoroids

Lastly, we used CCN to assess a relatively novel cancer model: tumoroids. We downloaded and assessed 131 distinct tumoroid expression profiles spanning 13 cancer categories from The NCI Patient-Derived Models Repository (PDMR)^69^ and from three individual studies^70–72^ (**Fig 7A, Supp Tab 7**). We note that several categories have three or fewer samples (BRCA, CESC, KIRP, OV, LIHC, and BLCA from PDMR). Among the cancer categories represented by more than three samples, only LUSC and PAAD have fewer than 50% classified as their annotated label (**Fig 7B**). In contrast to GBM CCLs, all three induced pluripotent stem cell-derived GBM tumoroids^72^ were classified as GBM with high CCN scores (mean = 0.53). To further characterize the tumoroids, we performed subtype classification on them (**Supp Tab 8**). UCEC tumoroids from PDMR contains a wide range of subtypes with two endometrioid, two serous and one mixed type (**Fig 7C**). On the other hand, LUSC tumoroids appear to be predominantly of classical subtypes with one tumoroid classified as a mix between classical and primitive (**Fig 7D**). Lastly, similar to the CCL and PDX counterparts, LUAD tumoroids are classified as proximal inflammatory and proximal proliferation with no tumoroids classified as TRU subtype (**Fig 7E**).

**Fig. 7.**
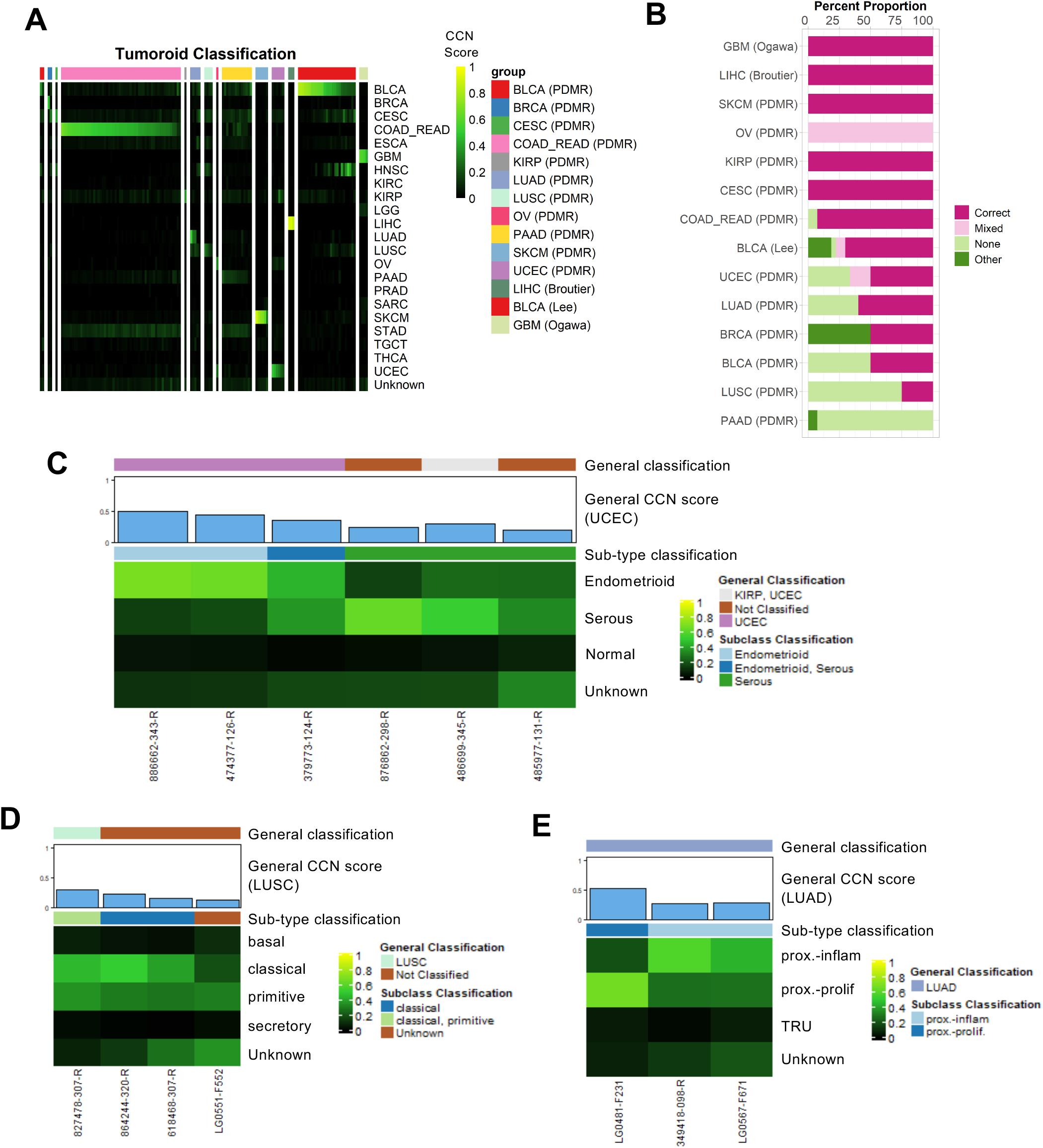
Evaluation of tumoroid models. **(A)** General classification heatmap of tumoroids. Column annotations represent annotated cancer type of the tumoroids, and row names represent cancer categories. **(B)** Proportion of classification categories in tumoroids across cancer types is visualized in the bar plot and ordered from the cancer type with highest proportion of correct and mixed correct classified tumoroids to the lowest. Subtype classification heatmap of **(C)** UCEC tumoroids, **(D)** LUSC tumoroids and **(E)** LUAD tumoroids. Only samples with CCN scores > 0.1 in their nominal tumor type are displayed.

### Comparison of CCLs, PDXs, GEMMs and tumoroids

Finally, we sought to estimate the comparative transcriptional fidelity of the four cancer models modalities. We compared the general CCN scores of each model on a per tumor type basis (**Fig 8**). In the case of GEMMs, we used the mean classification score of all samples with shared genotypes. We also used mean classification of technical replicates found in LIHC tumoroids^70^. We evaluated models based on both the maximum CCN score, as this represents the potential for a model class, and the median CCN score, as this indicates the current overall transcriptional fidelity of a model class. PDXs achieved the highest CCN scores in three (UCEC, PAAD, LUAD) out of the five cancer categories in which all four modalities were available (**Fig 8**), despite having low median CCN scores. Notably, PDXs have a median CCN score above the 0.25 threshold in PAAD while none of the other three modalities have any samples above the threshold. In LIHC, the highest CCN score for PDX (0.9) is only slightly lower than the highest CCN score for tumoroid (0.91). This suggest that certain individual PDXs most closely mimic the transcriptional state of native patient tumors despite a portion of the PDXs having low CCN scores. Similarly, while the majority of the CCLs have low CCN scores, several lines achieve high transcriptional fidelity in LUSC, LUAD and LIHC (**Fig 8**). Collectively, GEMMs and tumoroids had the highest median CCN scores in four of the five model classes (LUSC and LUAD for GEMMs and UCEC and LIHC for tumoroids). Notably, both of the LIHC tumoroids achieved CCN scores on par with patient tumors (**Fig 8**). In brief, this analysis indicates that PDXs and CCLs are heterogenous in terms of transcriptional fidelity, with a portion of the models highly mimicking native tumors and the majority of the models having low transcriptional fidelity (with the exception of PAAD for PDXs). On the other hand, GEMMs and tumoroids displayed a consistently high fidelity across different models.

**Fig. 8.**
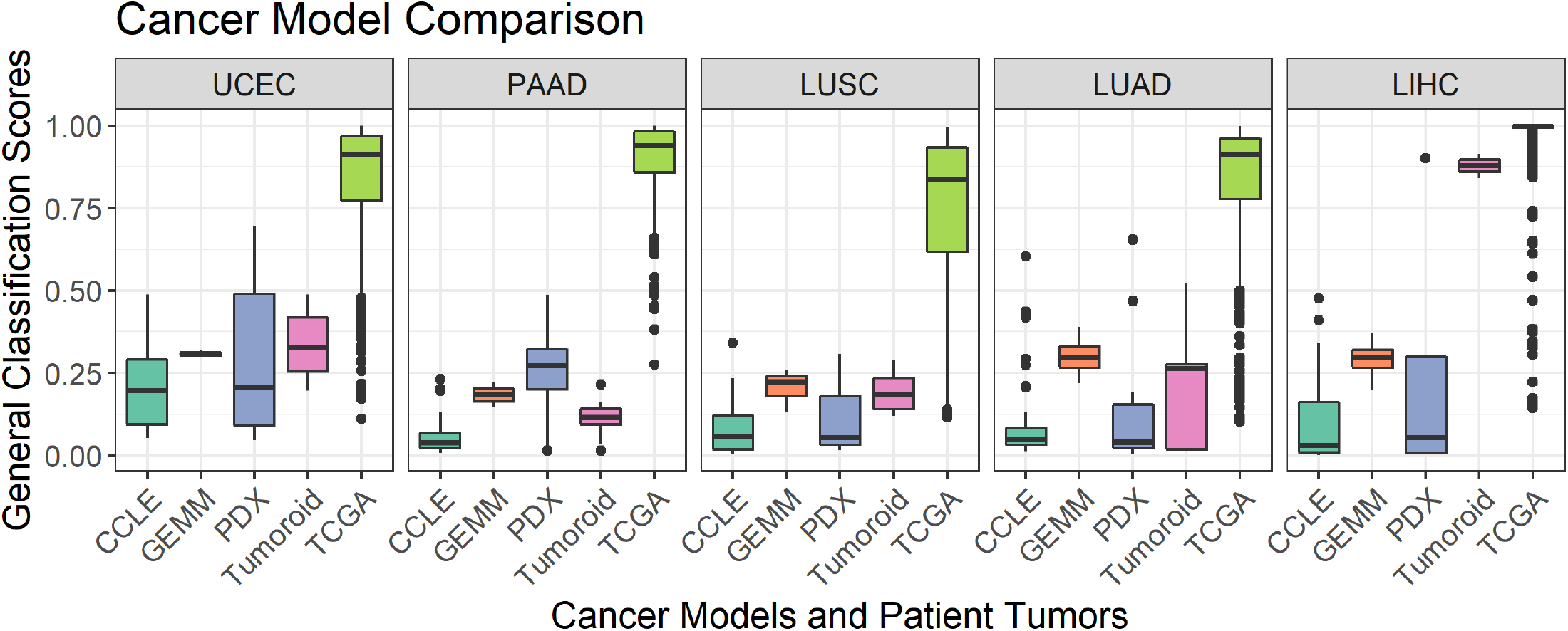
Comparison of CCLs, PDXs, and GEMMs. Box-and-whiskers plot comparing general CCN scores across CCLs, GEMMs, PDXs of five general tumor types (UCEC, PAAD, LUSC, LUAD, LIHC).

Because the CCN score is based on a moderate number of gene features (i.e. 1,979 gene pairs consisting of 1,689 unique genes) relative to the total number of protein-coding genes in the genome, it is possible that a cancer model with a high CCN score might not have a high global similarity to a naturally occurring tumor. Therefore, we also calculated the GRN status, a metric of the extent to which tumor-type specific gene regulatory network is established^21^, for all models (**Supp Fig 4**). We observed high level of correlation between the two similarity metrics, which suggests that although CCN classifies on a selected set of genes, its scores are highly correlated with global assessment of transcriptional similarity.

We also sought to compare model modalities in terms of the diversity of subtypes that they represent (**Supp Fig 5**). As a reference, we also included in this analysis the overall subtype incidence, as approximated by incidence in TCGA. Replicates in GEMMs and tumoroids were averaged into one classification profile. In models of UCEC, there is a notable difference in endometroid incidence, and the proportion of models classified as endometroid, with PDX and tumoroids having any representatives (**Supp Fig 5**). All of the CCL, GEMM, and tumoroid models of PAAD have an unknown subtype classification and no correct general classification. However, the majority of PDXs are subtyped as either a mixture of basal and classical, or classical alone. LUAD have proximal inflammation and proximal proliferation subtypes modelled by CCLs and PDX (**Supp Fig 5**). Likewise, LUSC have basal, classical and primitive subtypes modelled by CCLs and PDXs, and secretory subtype modelled by GEMMs exclusively (**Supp Fig 5**). Taken together, these results demonstrate the need to carefully select different model systems to more suitably model certain cancer subtypes.

## DISCUSSION

A major goal in the field of cancer biology is to develop models that mimic naturally occurring tumors with enough fidelity to enable therapeutic discoveries. However, methods to measure the extent to which cancer models resemble or diverge from native tumors are lacking. This is especially problematic now because there are many existing models from which to choose, and it has become easier to generate new models. Here, we present CancerCellNet (CCN), a computational tool that measures the similarity of cancer models to 22 naturally occurring tumor types and 36 subtypes. While the similarity of CCLs to patient tumors has already been explored in previous work, our tool introduces the capability to assess the transcriptional fidelity of PDXs, GEMMs, and tumoroids. Because CCN is platform- and species-agnostic, it represents a consistent platform to compare models across modalities including CCLs, PDXs, GEMMs and tumoroids. Here, we applied CCN to 657 cancer cell lines, 415 patient derived xenografts, 26 distinct genetically engineered mouse models and 131 tumoroids. Several insights emerged from our computational analyses that have implications for the field of cancer biology.

First, PDXs have the greatest potential to achieve transcriptional fidelity with three out of five general tumor types for which data from all modalities was available, as indicated by the high scores of individual PDXs. Notably PDXs are the only modality with samples classified as PAAD. At the same time, the median CCN scores of PDXs were lower than that of GEMMs and tumoroids in the other four tumor types. It is unclear what causes such a wide range of CCN scores within PDXs. We suspect that some PDXs might have undergone selective pressures in the host that distort the progression of genomic alterations away from what is observed in natural tumor^73^. Future work to understand this heterogeneity is important so as to yield consistently high fidelity PDXs, and to identify intrinsic and host-specific factors that so powerfully shape the PDX transcriptome.

Second, in general GEMMs and tumoroids have higher median CCN scores than those of PDXs and CCLs. This is also consistent with that fact that GEMMs are typically derived by recapitulating well-defined driver mutations of natural tumors, and thus this observation corroborates the importance of genetics in the etiology of cancer^74^. Moreover, in contrast to most PDXs, GEMMs are typically generated in immune replete hosts. Therefore, the higher overall fidelity of GEMMs may also be a result of the influence of a native immune system on GEMM tumors^75^. The high median CCN scores of tumoroids can be attributed to several factors including the increased mechanical stimuli and cell-cell interactions that come from 3D self-organizing cultures^76,77^.

Third, we have found that none of the samples that we evaluated here are transcriptionally adequate models of ESCA. This may be due to an inherent lability of the ESCA transcriptome that is often preceded by a metaplasia that has obscured determining its cell type(s) of origin^78^. Therefore, this tumor type requires further attention to derive new models.

Fourth, we found that in several tumor types, GEMMs tend to reflect mixtures of subtypes rather than conforming strongly to single subtypes. The reasons for this are not clear but it is possible that in the cases that we examined the histologically defined subtypes have a degree of plasticity that is exacerbated in the murine host environment.

Lastly, we recognize that many CCLs are not classified as their annotated labels. While we have suggested that the lack of immune component is not a major confounder, we suspect that the CCLs could undergo genetic divergence due to high number of passages, chemotherapy before biopsy, culture condition and genetic instability^79–82^, which could all be factors that drive CCLs away from their labelled tumors.

Currently, there are several limitations to our CCN tool, and caveats to our analyses which indicate areas for future work and improvement. First, CCN is based on transcriptomic data but other molecular readouts of tumor state, such as profiles of the proteome^83^, epigenome^84^, non-coding RNA-ome^84^, and genome^74^ would be equally, if not more important, to mimic in a model system. Therefore, it is possible that some models reflect tumor behavior well, and because this behavior is not well predicted by transcriptome alone, these models have lower CCN scores. To both measure the extent that such situations exist, and to correct for them, we plan in the future to incorporate other omic data into CCN so as to make more accurate and integrated model evaluation possible. As a first step in this direction, we plan to incorporate DNA methylation and genomic sequencing data as additional features for our Random forest classifier as this data is becoming more readily available for both training and cancer models. We expect that this will allow us to both refine our tumor subtype categories and it will enable more accurate predictions of how models respond to perturbations such as drug treatment.

A second limitation is that in the cross-species analysis, CCN implicitly assumes that homologs are functionally equivalent. The extent to which they are not functionally equivalent determines how confounded the CCN results will be. This possibility seems to be of limited consequence based on the high performance of the normal tissue cross-species classifier and based on the fact that GEMMs have the highest median CCN scores (in addition to tumoroids).

A third caveat to our analysis is that there were many fewer distinct GEMMs and tumoroids than CCLs and PDXs. As more transcriptional profiles for GEMMs and tumoroids emerge, this comparative analysis should be revisited to assess the generality of our results.

Finally, the TCGA training data is made up of RNA-Seq from bulk tumor samples, which necessarily includes non-tumor cells, whereas the CCLs are by definition cell lines of tumor origin. Therefore, CCLs theoretically could have artificially low CCN scores due to the presence of non-tumor cells in the training data. This problem appears to be limited as we found no correlation between tumor purity and CCN score in the CCLE samples. However, this problem is related to the question of intra-tumor heterogeneity. We demonstrated the feasibility of using CCN and single cell RNA-seq data to refine the evaluation of cancer cell lines contingent upon availability of scRNA-seq training data. As more training single cell RNA-seq data accrues, CCN would be able to not only evaluate models on a per cell type basis, but also based on cellular composition.

We have made the results of our analyses available online so that researchers can easily explore the performance of selected models or identify the best models for any of the 22 general tumor types and the 36 subtypes presented here. To ensure that CCN is widely available we have developed a free web application, which performs CCN analysis on user-uploaded data and allows for direct comparison of their data to the cancer models evaluated here. We have also made the CCN code freely available under an Open Source license and as an easily installed R package, and we are actively supporting its further development. Included in the web application are instructions for training CCN and reproducing our analysis. The documentation describes how to analyze models and compare the results to the panel of models that we evaluated here, thereby allowing researchers to immediately compare their models to the broader field in a comprehensive and standard fashion.

## Online Methods

### Training General CancerCellNet Classifier

To generate training data sets, we downloaded 8,991 patient tumor RNA-seq expression count matrix and their corresponding sample table across 22 different tumor types from TCGA using TCGAWorkflowData, TCGAbiolinks^85^ and SummarizedExperiment^86^ packages. We used all the patient tumor samples for training the general CCN classifier. We limited training and analysis of RNA-seq data to the 13,142 genes in common between the TCGA dataset and all the query samples (CCLs, PDXs, GEMMs, and tumoroids). To train the top pair Random forest classifier, we used a method similar to our previous method^23^. CCN first normalized the training counts matrix by down-sampling the counts to 500,000 counts per sample. To significantly reduce the execution time and memory of generating gene pairs for all possible genes, CCN then selected *n* up-regulated genes, *n* down-regulated genes and *n* least differentially expressed genes (CCN training parameter nTopGenes = *n*) for each of the 22 cancer categories using template matching^87^ as the genes to generate top scoring gene pairs. In short, for each tumor type, CCN defined a template vector that labelled the training tumor samples in cancer type of interest as 1 and all other tumor samples as 0 CCN then calculated the Pearson correlation coefficient between template vector and gene expressions for all genes. The genes with strong match to template as either upregulated or downregulated had large absolute Pearson correlation coefficient. CCN chose the upregulated, downregulated and least differentially expressed genes based on the magnitude of Pearson correlation coefficient.

After CCN selected the genes for each cancer type, CCN generated gene pairs among those genes. Gene pair transformation was a method inspired by the top-scoring pair classifier^88^ to allow compatibility of classifier with query expression profiles that were collected through different platforms (e.g. microarray query data applied to RNA-seq training data). In brief, the gene pair transformation compares 2 genes within an expression sample and encodes the “gene1_gene2” gene-pair as 1 if the first gene has higher expression than the second gene. Otherwise, gene pair transformation would encode the gene-pair as 0. Using all the gene pair combinations generated through the gene sets per cancer type, CCN then selected top *m* discriminative gene pairs (CCN training parameter nTopGenePairs = *m*) for each category using template matching (with large absolute Pearson correlation coefficient) described above. To prevent any single gene from dominating the gene pair list, we allowed each gene to appear at maximum of three times among the gene pairs selected as features per cancer type.

After the top discriminative gene pairs were selected for each cancer category, CCN grouped all the gene pairs together and gene pair transformed the training samples into a binary matrix with all the discriminative gene pairs as row names and all the training samples as column names. Using the binary gene pair matrix, CCN randomly shuffled the binary values across rows then across columns to generate random profiles that should not resemble training data from any of the cancer categories. CCN then sampled 70 random profiles, annotated them as “Unknown” and used them as training data for the “Unknown” category. Using gene pair binary training matrix, CCN constructed a multi-class Random Forest classifier of 2000 trees and used stratified sampling of 60 sample size to ensure balance of training data in constructing the decision trees.

To identify the best set of genes and gene-pair parameters (n and m), we used a grid-search cross-validation^89^ strategy with 5 cross-validations at each parameter set. The specific parameters for the final CCN classifier using the function “broadClass_train” in the package cancerCellNet are in **Supp Tab 9**. The gene-pairs are in **Supp Tab 10**.

### Validating General CancerCellNet Classifier

Two thirds of patient tumor data from each cancer type were randomly sampled as training data to construct a CCN classifier. Based on the training data, CCN selected the classification genes and gene-pairs and trained a classifier. After the classifier was built, 35 held-out samples from each cancer category were sampled and 40 “Unknown” profiles were generated for validation. The process of randomly sampling training set from 2/3 of all patient tumor data, selecting features based on the training set, training classifier and validating was repeated 50 times to have a more comprehensive assessment of the classifier trained with the optimal parameter set. To test the performance of final CCN on independent testing data, we applied it to 725 profiles from ICGC spanning 6 projects that do not overlap with TCGA (BRCA-KR, LIRI-JP, OV-AU, PACA-AU, PACA-CA, PRAD-FR).

### Selecting Decision Thresholds

Our strategy for selecting a decision threshold was to find the value that maximizes the average Macro F1 measure^90^ for each of the 50 cross-validations that were performed with the optimal parameter set, testing thresholds between 0 and 1 with a 0.01 increment. The F1 measure is defined as:

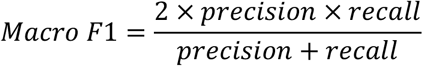

We selected the most commonly occurring threshold above 0.2 that maximized the average Macro F1 measure across the 50 cross-validations as the decision threshold for the final classifier (threshold = 0.25). The same approach was applied for the subtype classifiers. The thresholds and the corresponding average precision, recall and F1 measures are recorded in (**Supp Tab 11**).

### Classifying Query Data into General Cancer Categories

We downloaded the RNA-seq cancer cell lines expression profiles and sample table from (https://portals.broadinstitute.org/ccle/data), and microarray cancer cell lines expression profiles and sample table from Barretina et al^37^. We extracted two WT control NCCIT RNA-seq expression profiles from Grow et al^91^. We received PDX expression estimates and sample annotations from the authors of Gao et al ^20^. We gathered GEMM expression profiles from nine different studies^59–67^. We downloaded tumoroid expression profiles from The NCI Patient-Derived Models Repository (PDMR)^69^ and from three individual studies^70–72^. To use CCN classifier on GEMM data, the mouse genes from GEMM expression profiles were converted into their human homologs. The query samples were classified using the final CCN classifier. Each query classification profile was labelled as one of the four classification categories: “correct”, “mixed”, “none” and “other” based on classification profiles. If a sample has a CCN score higher than the decision threshold in the labelled cancer category, we assigned that as “correct”. If a sample has CCN score higher than the decision threshold in labelled cancer category and in other cancer categories, we assigned that as “mixed”. If a sample has no CCN score higher than the decision threshold in any cancer category or has the highest CCN score in ‘Unknown’ category, then we assigned it as “none”. If a sample has CCN score higher than the decision threshold in a cancer category or categories not including the labelled cancer category, we assigned it as “other”. We analyzed and visualized the results using R and R packages pheatmap^92^ and ggplot2^93^.

### Cross-Species Assessment

To assess the performance of cross-species classification, we downloaded 1003 labelled human tissue/cell type and 1993 labelled mouse tissue/cell type RNA-seq expression profiles from Github (https://github.com/pcahan1/CellNet). We first converted the mouse genes into human homologous genes. Then we found the intersecting genes between mouse tissue/cell expression profiles and human tissue/cell expression profiles. Limiting the input of human tissue RNA-seq profiles to the intersecting genes, we trained a CCN classifier with all the human tissue/cell expression profiles. The parameters used for the function “broadClass_train” in the package cancerCellNet are in **Supp Tab 9**. We randomly sampled 75 samples from each tissue category in mouse tissue/cell data and applied the classifier on those samples to assess performance.

### Cross-Technology Assessment

To assess the performance of CCN in applications to microarray data, we gathered 6,219 patient tumor microarray profiles across 12 different cancer types from more than 100 different projects (**Supp Tab 12**). We found the intersecting genes between the microarray profiles and TCGA patient RNA-seq profiles. Limiting the input of RNA-seq profiles to the intersecting genes, we created a CCN classifier with all the TCGA patient profiles using parameters for the function “broadClass_train” listed in **Supp Tab 9**. After the microarray specific classifier was trained, we randomly sampled 60 microarray patient samples from each cancer category and applied CCN classifier on them as assessment of the cross-technology performance in **Supp Fig 2A**. The same CCN classifier was used to assess microarray CCL samples **Supp Fig 2B**.

### Training and validating scRNA-seq Classifier

We extracted labelled human melanoma and glioblastoma scRNA-seq expression profiles^40,41^, and compiled the two datasets excluding 3 cell types T.CD4, T.CD8 and Myeloid due to low number of cells for training. 60 cells from each of the 11 cell types were sampled for training a scRNA-seq classifier. The parameters for training a general scRNA-seq classifier using the function “broadClass_train” are in **Supp Tab 9.** 25 cells from each of the 11 cell types from the held-out data were selected to assess the single cell classifier. Using maximization of average Macro F1 measure, we selected the decision threshold of 0.255. The gene-pairs that were selected to construct the classifier are in **Supp Tab 10**. To assess the cross-technology capability of applying scRNA-seq classifier to bulk RNA-seq, we downloaded 305 expression profiles spanning 4 purified cell types (B cells, endothelial cells, monocyte/macrophage, fibroblast) from https://github.com/pcahan1/CellNet.

### Training Subtype CancerCellNet

We found 11 cancer types (BRCA, COAD, ESCA, HNSC, KIRC, LGG, PAAD, UCEC, STAD, LUAD, LUSC) which have meaningful subtypes based on either histology or molecular profile and have sufficient samples to train a subtype classifier with high AUPR. We also included normal tissues samples from BRCA, COAD, HNSC, KIRC, UCEC to create a normal tissue category in the construction of their subtype classifiers. Training samples were either labelled as a cancer subtype for the cancer of interest or as “Unknown” if they belong to other cancer types. Similar to general classifier training, CCN performed gene pair transformation and selected the most discriminate gene pairs for each cancer subtype. In addition to the gene pairs selected to discriminate cancer subtypes, CCN also performed general classification of all training data and appended the classification profiles of training data with gene pair binary matrix as additional features. The reason behind using general classification profile as additional features is that many general cancer types may share similar subtypes, and general classification profile could be important features to discriminate the general cancer type of interest from other cancer types before performing finer subtype classification. The specific parameters used to train individual subtype classifiers using “subClass_train” function of CancerCellNet package can be found in **Supp Tab 9** and the gene pairs are in **Supp Tab 10**.

### Validating Subtype CancerCellNet

Similar to validating general class classifier, we randomly sampled 2/3 of all samples in each cancer subtype as training data and sampled an equal amount across subtypes in the 1/3 held-out data for assessing subtype classifiers. We repeated the process 20 times for more comprehensive assessment of subtype classifiers.

### Classifying Query Data into Subtypes

We assigned subtype to query sample if the query sample has CCN score higher than the decision threshold. The table of decision threshold for subtype classifiers are in **Supp Tab 11**. If no CCN scores exceed the decision threshold in any subtype or if the highest CCN score is in ‘Unknown’ category, then we assigned that sample as ‘Unknown’. Analysis was performed in R and visualizations were generated with the ComplexHeatmap package^94^.

### Cells culture, Immunohistochemistry and histomorphometry

Caov-4 (ATCC^®^ HTB-76™), SK-OV-3(ATCC^®^ HTB-77™), RT4 (ATCC^®^ HTB-2™), and NCCIT(ATCC^®^ CRL-2073™) cell lines were purchased from ATCC. HEC-59 (C0026001) and A2780 (93112519-1VL) were obtained from Addexbio Technologies and Sigma-Aldrich. Vcap and PC-3. SK-OV-3, Vcap, and RT4 were cultured in Dulbecco’s Modified Eagle Medium (DMEM, high glucose, 11960069, Gibco) with 1% Penicillin-Streptomycin-Glutamine (10378016, Life Technologies); Caov-4, PC-3, NCCIT, and A2780 were cultured using RPMI-1640 medium (11875093, Gibco) while HEC-59 was in Iscove’s Modified Dulbecco’s Medium (IMDM, 12440053, Gibco). Both media were supplemented with 1% Penicillin-Streptomycin (15140122, Gibco). All medium included 10% Fetal Bovine Serum (FBS).

Cells cultured in 48-well plate were washed twice with PBS and fixed in 10% buffered formalin for 24 hrs at 4 °C. Immunostaining was performed using a standard protocol. Cells were incubated with primary antibodies to goat HOXB6 (10 μg/mL, PA5-37867, Invitrogen), mouse WT1(10 μg/mL, MA1-46028, Invitrogen), rabbit PPARG (1:50, ABN1445, Millipore), mouse FOLH1(10 μg/mL, UM570025, Origene), and rabbit LIN28A (1:50, #3978, Cell Signaling) in Antibody Diluent (S080981-2, DAKO), at 4 °C overnight followed with three 5 min washes in TBST. The slides were then incubated with secondary antibodies conjugated with fluorescence at room temperature for 1 h while avoiding light followed with three 5 min washes in TBST and nuclear stained with mounting medium containing DAPI. Images were captured by Nikon EcLipse Ti-S, DS-U3 and DS-Qi2.

Histomorphometry was performed using ImageJ (Version 2.0.0-rc-69/1.52i). % N.positive cells was calculated by the percentage of the number of positive stained cells divided by the number of DAPI-positive nucleus within three of randomly chosen areas. The data were expressed as means ± SD.

### Tumor Purity Analysis

We used the R package ESTIMATE^95^ to calculate the ESTIMATE scores from TCGA tumor expression profiles that we used as training data for CCN classifier. To calculate tumor purity we used the equation described in YoshiHara et al., 2013^95^:

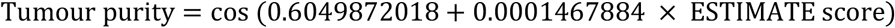

### Extracting Citation Counts

We used the R package RISmed^96^ to extract the number of citations for each cell line through query search of “*cell line name*[Text Word] AND *cancer*[Text Word]” on PubMed. The citation counts were normalized by dividing the citation counts with the number of years since first documented.

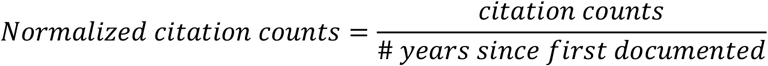

### GRN construction and GRN Status

GRN construction was extended from our previous method^21^. 80 samples per cancer type were randomly sampled and normalized through down sampling as training data for the CLR GRN construction algorithm. Cancer type specific GRNs were identified by determining the differentially expressed genes per each cancer type and extracting the subnetwork using those genes.

To extend the original GRN status algorithm^21^ across different platforms and species, we devised a rank-based GRN status algorithm. Like the original GRN status, rank based GRN status is a metric of assessing the similarity of cancer type specific GRN between training data in the cancer type of interest and query samples. Hence, high GRN status represents high level of establishment or similarity of the cancer specific GRN in the query sample compared to those of the training data. The expression profiles of training data and query data were transformed into rank expression profiles by replacing the expression values with the rank of the expression values within a sample (highest expressed gene would have the highest rank and lowest expressed genes would have a rank of 1). Cancer type specific mean and standard deviation of every gene’s rank expression were learned from training data. The modified Z-score values for genes within cancer type specific GRN were calculated for query sample’s rank expression profiles to quantify how dissimilar the expression values of genes in query sample’s cancer type specific GRN compared to those of the reference training data:

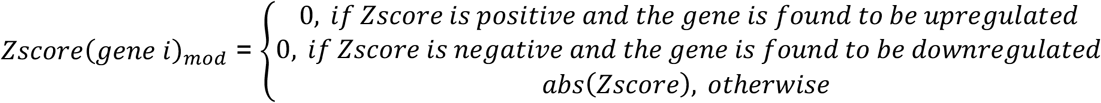

If a gene in the cancer type specific GRN is found to be upregulated in the specific cancer type relative to other cancer types, then we would consider query sample’s gene to be similar if the ranking of the query sample’s gene is equal to or greater than the mean ranking of the gene in training sample. As a result of similarity, we assign that gene of a Z-score of 0. The same principle applies to cases where the gene is downregulated in cancer specific subnetwork.

GRN status for query sample is calculated as the weighted mean of the (1000 – *Zscore*(*gene i*)_*mod*_) across genes in cancer type specific GRN. 1000 is an arbitrary large number, and larger dissimilarity between query’s cancer type specific GRN indicate high Z-scores for the GRN genes and low GRN status.

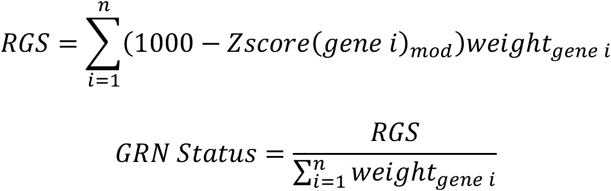

The weight of individual genes in the cancer specific network is determined by the importance of the gene in the Random Forest classifier. Finally, the GRN status gets normalized with respect to the GRN status of the cancer type of interest and the cancer type with the lowest mean GRN status.

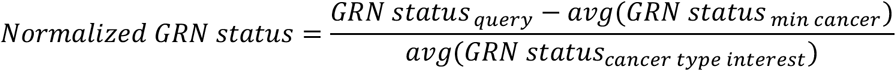

Where “min cancer” represents the cancer type where its training data have the lowest mean GRN status in the cancer type of interest, and *avg*(*GRN status_min cancer_*) represents the lowest average GRN status in the cancer type of interest. *avg*(*GRN status_cancer type interest_*) represents average GRN status of the cancer type of interest in the training data.

### Code availability

CancerCellNet code and documentation is available at GitHub: https://github.com/pcahan1/cancerCellNet

## Acknowledgements

This work was supported by the National Institutes of Health NCI Ovarian Cancer SPORE P50CA228991 via a Development Research Program award to PC. FWH was supported by a Prostate Cancer Foundation Young Investigator Award, Department of Defense W81XWH-17-PCRP-HD (F.W.H.), the National Institutes of Health/National Cancer Institute P20 CA233255-01 (F.W.H.) U19 CA214253 (F.W.H.). We would like to thank John Powers, Hao Zhu, Tian-Li Wang, Charles Eberhart, and Kaloyan Tsanov for comments on the manuscript and helpful discussions. Some figures were created in part with Biorender.com.

## Supplementary Information

**Supplementary Figure 1.**
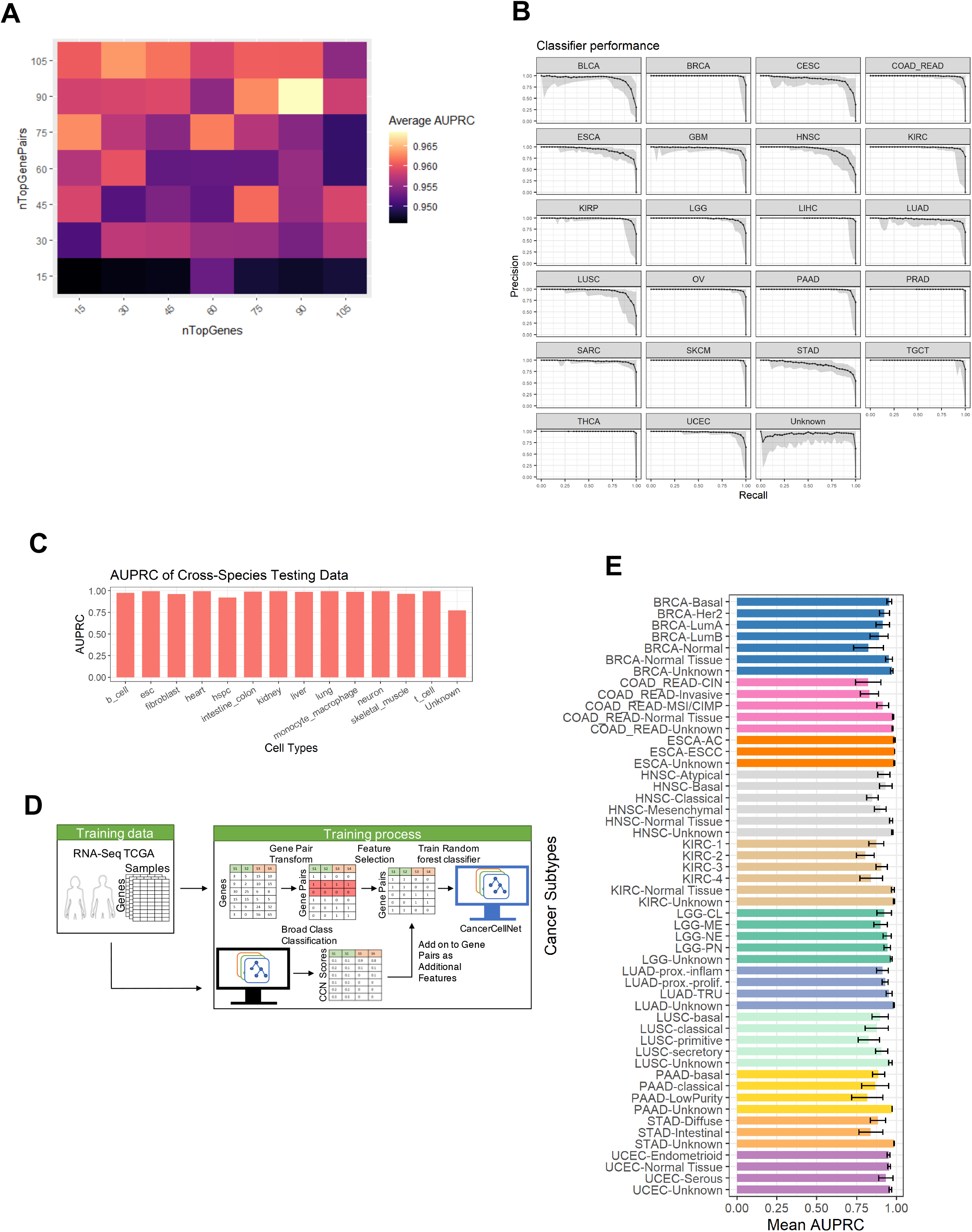
Assessment of CCN general classifier and subtype classifier. **(A)** Mean AUPRC of repeated grid-search cross-validation for each parameter grid. **(B)** Mean and range of CCN classifier’s PR curves from 50 cross validations based on the optimal feature selection parameters *n* and *m*. **(C)** AUPRC of CCN human tissue classifier when applied to mouse tissue data. **(D)** The schematic of training a subtype classifier in CCN. CCN uses patient tumor expression profiles from cancer of interest as training data. CCN performs gene-pair transformation and selects the most discriminative gene pairs among the cancer subtypes from training data as features. CCN then applies the general classification on training data and uses the general classification profile as features in addition to gene pairs for training a Random Forest classifier. The weight of the general classification profiles as features can be tuned to improve AUPRC. **(E)** The mean and standard deviation of AUPRC for 11 subtype classifiers based on 20 iterations of random sampling of training and held-out data, training subtype classifier using training data, classification of held-out data, and calculation of recall and precision.

**Supplementary Figure 2.**
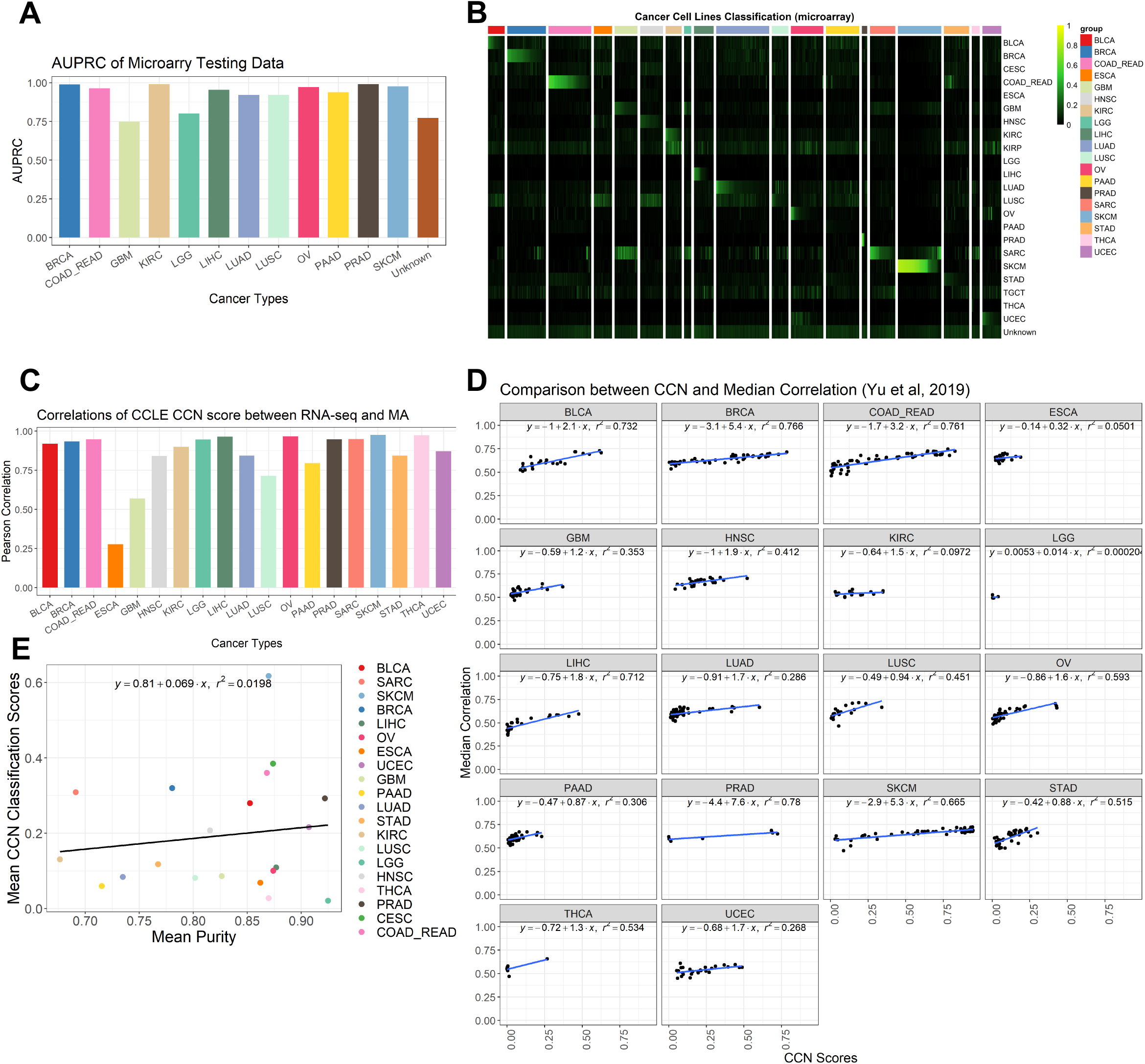
Further validation of CCN and classification results. To validate the cross-platform classification performance of CCN, a new classifier specifically trained to classify microarray data was trained using RNA-seq data from TCGA as training data and intersecting genes between RNA-seq data and microarray data. **(A)** AUPRC of CCN classifier when applied to tumor profiles assayed on microarrays. **(B)** Classification heatmap of CCLs using microarray expression data. **(C)** Pearson correlation between CCN scores of CCLE lines generated from RNA-seq data and microarray data. **(D)** Comparison between CCLs’ CCN scores and the similarity metric from Yu et al^15^, median correlations of transcriptional profiles between CCLs and TCGA tumors from CCLs’ labelled cancer category. **(E)** Comparison of mean tumor purity of training data and mean CCN scores of CCLs for each cancer category.

**Supplementary Figure 3.**
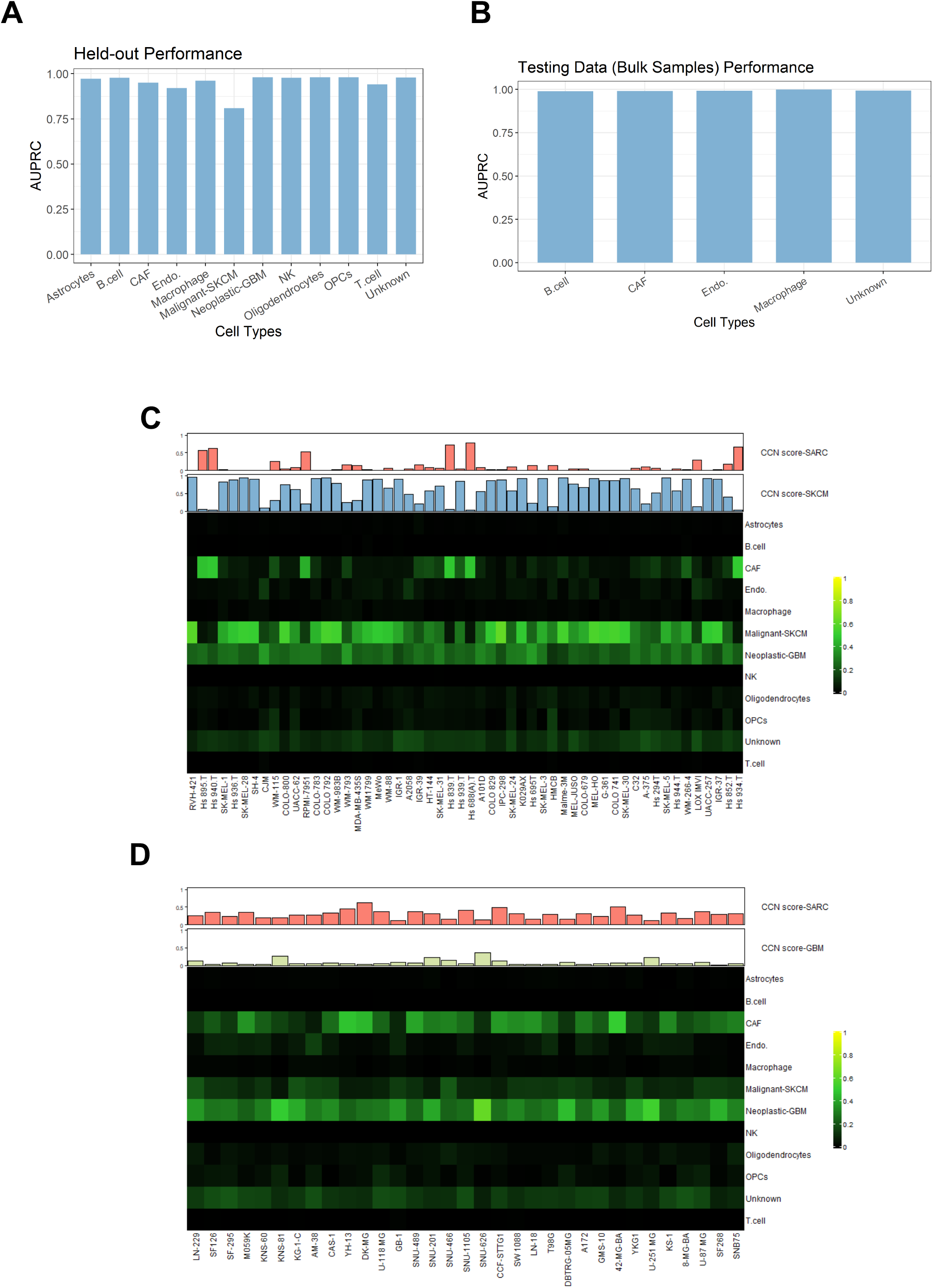
Single-cell classification of SKCM and GBM cell lines. **(A)** AUPRC of the single-cell classifier when applied to scRNA-seq held-out data. **(B)** AUPRC of the scRNA-seq classifier when applied to purified bulk RNA samples. **(C)** Single-cell classification of SKCM CCLs. Red bar-plot (top) represents general CCN scores in SARC and blue bar-plot (bottom) represents general CCN scores in SKCM. **(D)** Single-cell classification of GBM CCLs. Red bar-plot (top) represents general CCN scores in SARC and yellow bar-plot (bottom) represents general CCN scores in GBM.

**Supplementary Figure 4.**
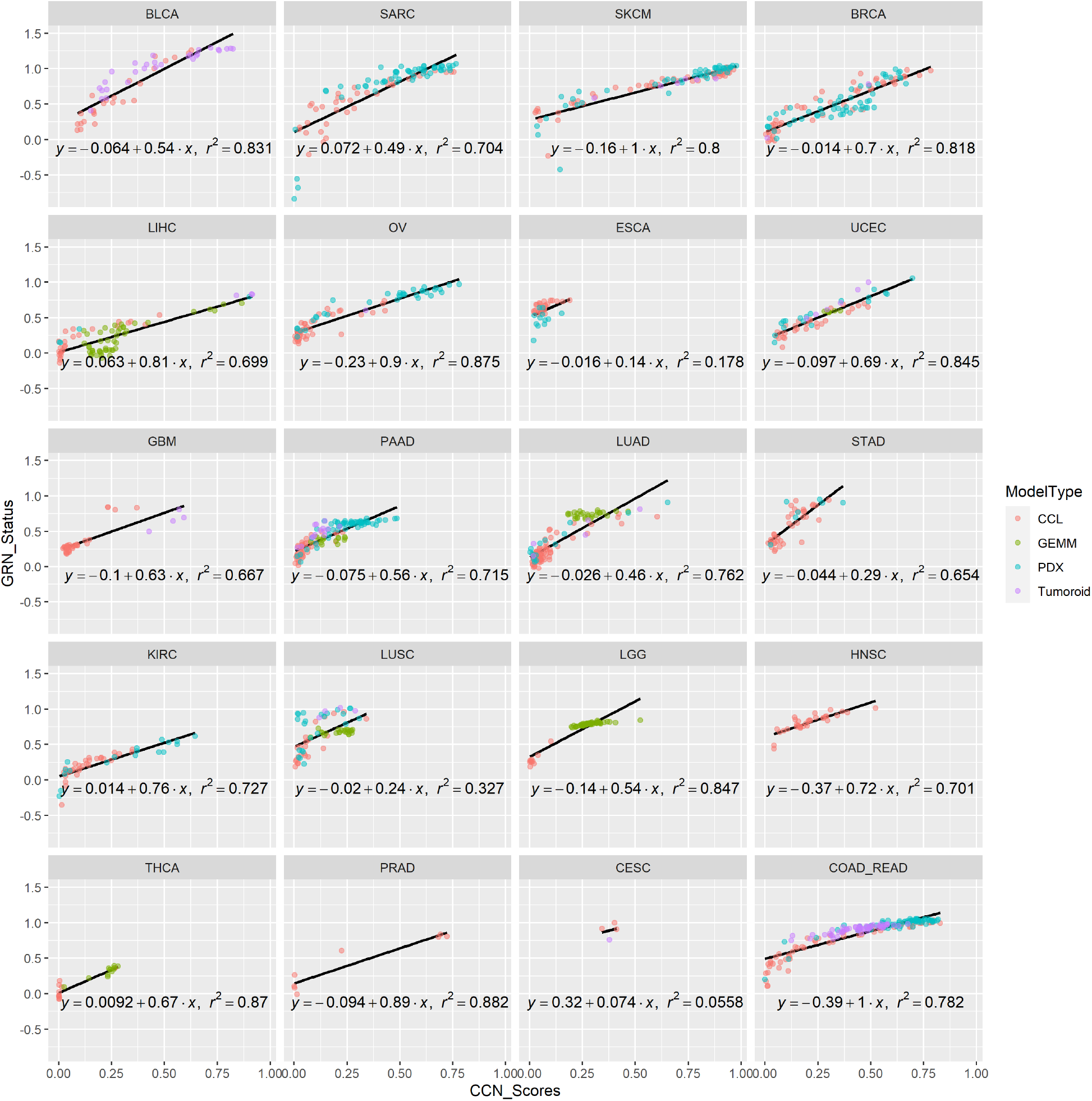
Correlation between cancer type specific network GRN status and general CCN scores.

**Supplementary Figure 5.**
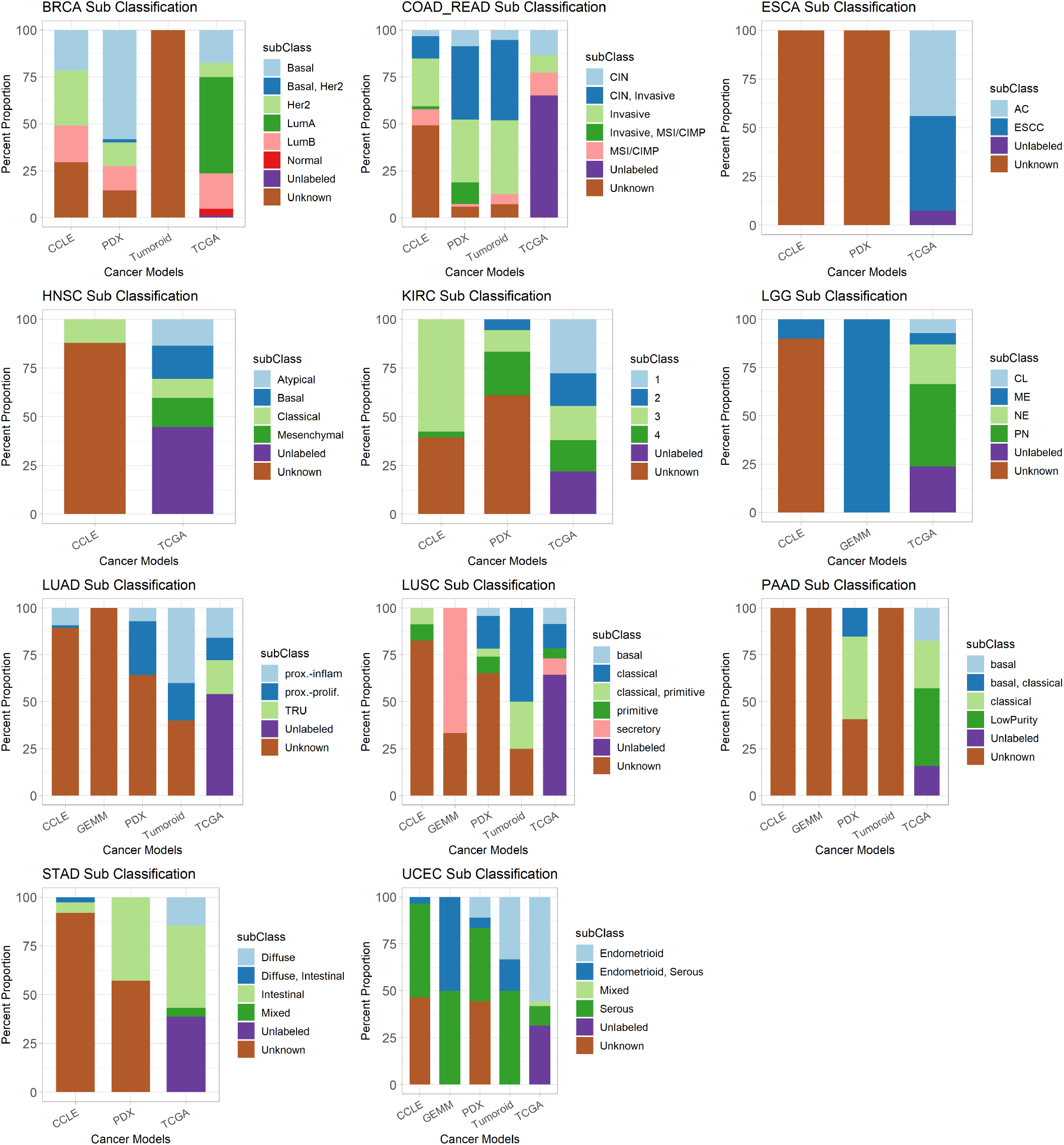
Proportion of cancer subtypes in different cancer models and TCGA tumor data across 11 general cancer types.

**Supplementary Table 1** General classification profiles of CCLs.

**Supplementary Table 2** Subtype classification profiles of CCLs.

**Supplementary Table 3** General classification profiles of PDXs.

**Supplementary Table 4** Subtype classification profiles of PDXs.

**Supplementary Table 5** General classification profiles of GEMMs

**Supplementary Table 6** Subtype classification profiles of GEMMs.

**Supplementary Table 7** General classification profiles of tumoroids.

**Supplementary Table 8** Subtype classification profiles of tumoroids.

**Supplementary Table 9** Specific parameters used for training of all classifiers.

**Supplementary Table 10** Gene-pairs selected for final training of CCN general, subtype classifiers and single-cell classifier.

**Supplementary Table 11** Decision thresholds and the corresponding precision and recall for the general classifier and subtype classifier.

**Supplementary Table 12** Accessions of tumor microarray data used in validation.

